# Matrix stiffness and blood pressure together regulate vascular smooth muscle cell phenotype switching

**DOI:** 10.1101/2020.12.27.424498

**Authors:** Pamela Swiatlowska, Brian Sit, Zhen Feng, Emilie Marhuenda, Ioannis Xanthis, Simona Zingaro, Matthew Ward, Xinmiao Zhou, Qingzhong Xiao, Cathy Shanahan, Gareth E Jones, Cheng-han Yu, Thomas Iskratsch

**Author notes:** Authors contributed equally (Pamela Swiatlowska, Brian Sit, Zhen Feng). Correspondence: Thomas Iskratsch, Cheng-han Yu.

## Abstract

Vascular smooth muscle cells (VSMCs) play a central role in the onset and progression of atherosclerosis. In pre-atherosclerotic lesions, VSMCs switch from a contractile to a synthetic phenotype and subsequently remodel the microenvironment, leading to further disease progression. Ageing and associated mechanical changes of the extracellular matrix as well as hypertension are major risk of atherosclerosis. Consequently, we sought here to systematically study the impact of mechanical stimulations on VSMC phenotypic switching, by modulating both stiffness and hydrodynamic pressure. Thereby we find that hemodynamic pressure and matrix stiffness individually affect the VSMC phenotype. However, only the combination of hypertensive pressure and matrix compliance, and as such mechanical stimuli that are prevalent during atherosclerosis, lead to a full phenotypic switch including the formation of matrix degrading podosomes. We further analyse the molecular mechanism in stiffness and pressure sensing and identify a regulation through different, but overlapping pathways, culminating in the regulation of the actin cytoskeleton through cofilin. Altogether, our data shows how different pathological mechanical signals combined, but through distinct pathways accelerate a phenotypic switch that will ultimately contribute to atherosclerotic disease progression.

## Introduction

Cardiovascular diseases are the primary cause of mortality worldwide. Vascular ageing is dramatically increasing the risk of cardiovascular disease, including systolic hypertension, coronary artery disease, stroke, heart failure and atrial fibrillation or atherosclerosis(*1–5*). This ageing process is related to both chemical/molecular as well as mechanical changes, including changes to the blood flow (wall stress, haemodynamic pressure) and stiffness of the arterial wall. Endothelial cells (ECs) sense the wall stress and shear direction, and the associated mechanosignalling defines atheroprone or atheroprotective regions(*6*). The shear stress deforms the apical surface of ECs and is sensed amongst others by the glycocalyx, mechanosensitive ion channels and various receptors including PECAM-1, plexin D1, and integrins(*6–10*). ECs communicate this then further to the underlying arterial layers and the residing vascular smooth muscle cells (VSMCs), through the extracellular matrix, growth factor signalling and direct receptor interaction(*11*).

VSMCs are necessary for matrix formation and contractility of the arterial wall but can adopt alternative phenotypes in response to chemical and physical signals. The phenotypically altered cells have been described as synthetic, macrophage-like, or foam cells(*12*, *13*). Previous studies on the involvement of VSMCs in atherosclerosis have largely focused on their role in advanced atherosclerotic processes i.e., fibrous cap formation and plaque stability or rupture. However recent research increasingly points to a critical role for VSMCs in the initiation and early phases of atherosclerosis, including diffuse intimal thickenings (DIT) - the most likely precursor of pre-atherosclerotic plaques (*12*, *14*), as well as pathological intimal thickening (PIT), which is considered the first stage of atherosclerosis (*12*). Moreover, lineage tracing data shed light into origin of the atherosclerotic VSMCs and identified that especially the VSMC within the intima layer change their phenotype to define the disease onset and progression (*12*, *15*, *16*). In DITs, VSMCs are considered the major source of extracellular matrix proteins and especially proteoglycans (versican and biglycan, as well as others to a lesser degree)(*17*) in the arterial wall, hence leading to the thickening of the intima and the progression to PIT (*12*, *18*). This progression is promoted by retention of apolipoproteins, oxidation of lipids, VSMC phenotypic switching, proliferation and apoptosis. Again, the capability of (phenotypically altered) VSMCs to secrete and remodel the extracellular matrix is critical for this step(*19–22*).

In addition to the role of EC-VSMC cross-signalling, there is circumstantial evidence suggesting that mechanical forces also direct impact on the VSMC phenotype. DIT has long been considered an adaptation to blood flow (*23*). Moreover, mechanical strain altered mRNA expression and accumulation of extracellular matrix proteins from VSMCs (*22*). Finally, pressure induced phenotypic changes and migration in VSMCs (*24*, *25*) - although these studies used static pressure and rigid tissue culture plastic and therefore non-physiological stimuli.

Arteries are composite structures with complex mechanical behaviour. The mechanical properties differ between macro and microscale, with micro-scale stiffnesses magnitudes lower compared to the macroscale (*26–30*). Further, comparative studies between the individual arterial wall layers suggested that the intima was more compliant compared to the media layer (~5kPa vs ~ 40kPa measured in mouse ascending aortas)(*31*). Also, measurements of lipid rich regions in (human and ApoE−/− mouse) atherosclerotic plaques respectively indicated a compliant environment (2.7 ± 1.8 kPa and 5.5 ± 3.5 kPa, respectively) and potentially further softening at the microscale during DIT and PIT where proteoglycan and lipid content increases (*12*, *18*, *29*, *32*).

The expression of proteoglycans not only affects the compliance but also results in a larger proportion of void space and hence, increased compressibility of the intima and the residing VSMCs (which have been described as poro-elastic due to the expression of aquaporins) (*28*, *29*, *33–41*). Conclusively, previous studies suggested that the compression of the intima increases (*36*), while its stretching decreases due to the accompanying arterial macroscale stiffening (*26*, *27*, *42–46*).

Because of the changing mechanical stimuli in ageing and vascular disease, we sought here to study the effects of the different mechanical stimuli (pressure, stretch, rigidity) in isolation, to identify critical parameters influencing phenotypic changes of the VSMCs. Thereby, we find individual effects of pressure and compliance which both favour a phenotypic switch. This switch is potentiated when combining compliance and pressure. In combination of these two stimuli, we find large-scale changes to protein expression, cell morphology, actin organisation, and maximal formation of matrix degrading podosomes. We further study the regulation of the podosomes downstream of mechanical signals and identify distinct molecular pathways, both acting on the same protein, cofilin, which regulates the podosome formation and turnover. Together our data highlights the strong contribution of different mechanical factors on VSMCs that in combination will influence atherosclerotic disease progression.

## Results

### The phenotypic switch of vascular smooth muscle cells is regulated through haemodynamic pressure and matrix stiffness

Vascular smooth muscle cells (VSMCs) play a key role in various stages of atherosclerosis and disease progression is associated with phenotypic switching. Extracellular matrix changes and hypertensive blood pressure are both critically linked to the onset and progression of atherosclerosis. Therefore, we wanted to study the effect the different mechanical stimuli (rigidity, pressure, stretch) on vascular smooth muscle cell phenotype. Specifically, we plated A7r5 VSMCs onto PDMS coated coverslips with defined stiffness covering the whole range that VSMCs are exposed to, from an intima with enhanced proteoglycan expression and lipid inclusions (1kPa) to healthy media layer (20kPa) to and calcified regions in the atherosclerotic plaques (130kPa). In addition, the VSMCs on the stiffness-defined substrates were subjected to hydrodynamic pressure stimulation, mimicking either healthy normal blood pressure (NBP, 120/60 mmHg) or stage II hypertension (HT, 180/120 mmHg) (Fig 1A), using a pressure stimulator (CellScale Mechanoculture TR, modified for a low-pressure range), situated inside a 37°C tissue culture incubator. Control cells were kept inside the chamber without the pressure stimulation. After 12 hours, cells were fixed and stained with phalloidin before cell segmentation and analysis of morphological parameters with CellProfiler (cell shape and F-actin shape descriptors) and the ImageJ OrientationJ plugin (F-actin alignment)(*47*, *48*). To detect F-actin dots, which we initially observed in some cells, we trained a pixel classification using Ilastik and measured the number and the area of dots in cells after the segmentation (Fig 1B). We then performed a dimensionality reduction using t-distributed Stochastic Neighbor Embedding (t-SNE) (Fig 1C), and two separate clusters were identified, including primary control and NBP treated cells (cluster 1) or HT treated cells (cluster 2) (Fig 1D). Overall, we noted a correlation of the cluster composition with the pressure stimulation regime and, to a smaller degree, with the substrate stiffness (Fig 1D). Consistent with this, cell shape analysis using the visually-aided morpho-phenotyping image recognition “VAMPIRE” software(*49*) indicated an increase in round, compared to spindle shaped cells after the HT treatment (Fig 1E), while NBP treatment in contrast preserved spindle shapes and further resulted in an apparent increase in cellular alignment (not shown). After HT treatment, we also found a substantial change to the actin organisation, including a reduction in actin alignment and overall organisation (Fig 1F). Specifically, we also detected changes to actin dot occurrence, which occurred either as single dots or clusters (Fig 1G).

**Figure 1:**
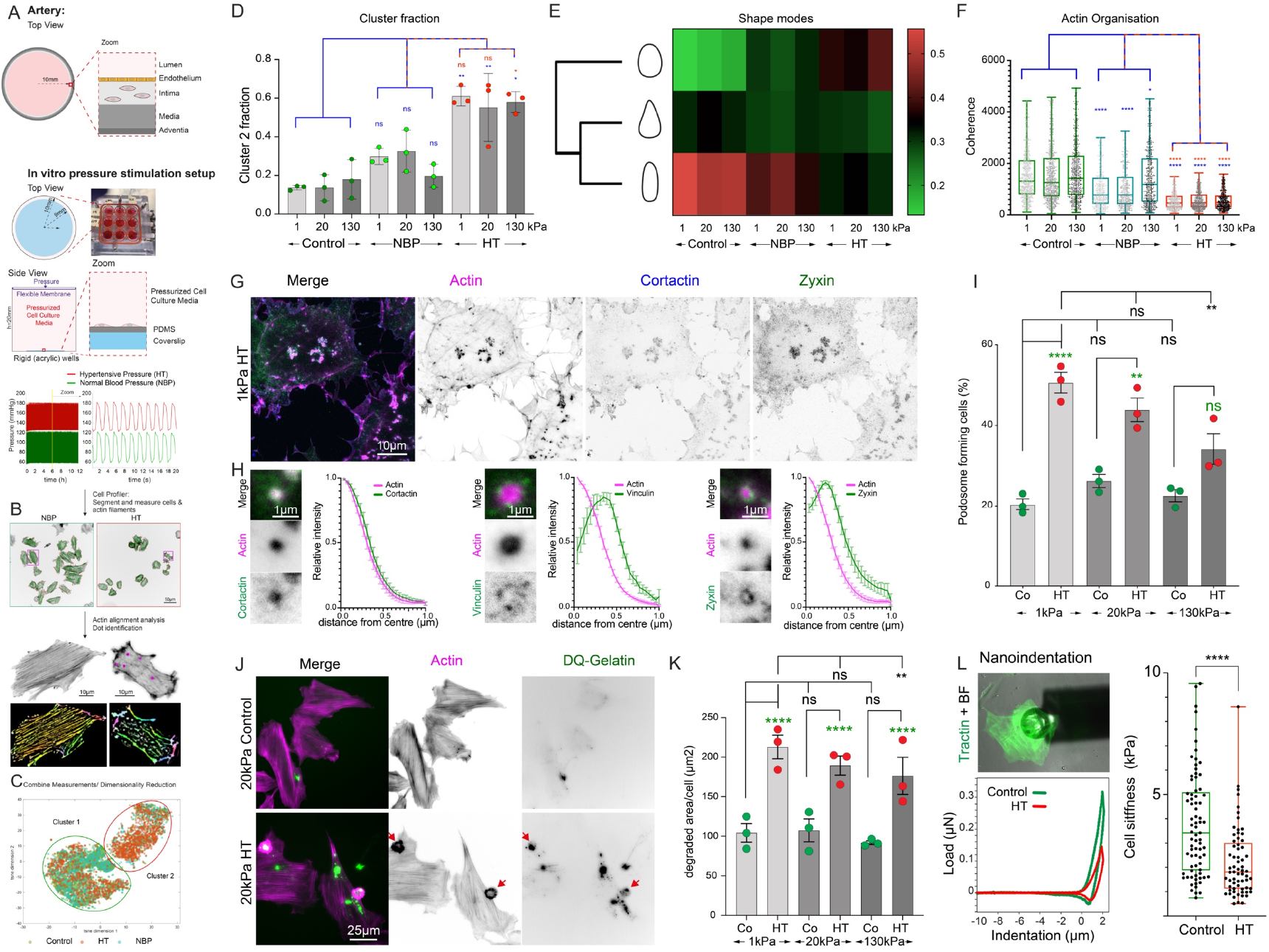
Hypertensive pressure, but not healthy blood pressure stimulates phenotypic switching in vascular smooth muscle cells. A) overview of the experimental protocol. Cells were plated on PDMS coated coverslips with different stiffness (1,20,130kPa) and placed under cyclic hydrodynamic pressure mimicking either healthy (120/60mmHg) or hypertensive blood pressure (180/120mmHg); measured output from stimulator shown on left, zoom into region denoted by yellow line shown on top right. B) Cells were then fixed, stained and analysed using Cell Profiler for cell (middle panel, red outlines) and actin segmentation (green outlines). Segmented cells were further analysed to determine actin dots (purple outlines) or actin filament alignment (bottom panel). C) Area, shape, actin shape, orientation and actin dot measurements were combined before dimensionality reduction and cluster analysis (t-SNE) D) The presence of the various conditions in each cluster was analysed after separation of the data from the three different repeats. Error bars: SD; E) HT treatment affected shape modes of the cells. F) HT treatment resulted in reduced actin organisation (Coherence measurements using the orientationJ plugin, pooled data from three separate experiments) G) Actin dots appeared both in single dots and clusters and co-stained for cortactin, zyxin or vinculin (H), with zyxin and vinculin appearing in a ring-structure around the central actin core when imaged with super-resolution spinning disc microscopy (H); radial profiles are normalized intensities from n=10 podosomes from 5 cells each. I) The fraction of podosome-forming cells depended on pressure and stiffness (fraction of podosome-forming cells, displayed as average per repeat, from three independent experiments with 10 images each and ~10-20 cells per image). J,K) Podosomes actively degraded gelatin, but the efficiency depended on stiffness and pressure (DQ-gelatin positive area per cell, displayed as average per repeat, from n=200-300 cells per repeat and condition). L) the large-scale changes to the cellular cytoskeleton were also reflected by a drop in the cellular stiffness as determined by nanoindentation (data from three independent experiments). Top left: image from tractin (green) and the indenter (bright field channel). Bottom left: typical load-indentation curve for the control (green) and HT (red) treated cells. Right panel: Young’s moduli. *p < 0.0332; **p < 0.0021; ***p < 0.0002; ****p < 0.0001; ns, not significant; p-values from ANOVA (F), and Tukey test for multiple comparisons or unpaired t-test (L); D,F: blue asterisks are for comparison with control, red asterisks for comparison with NBP; I,K: green asterisks represent comparison to the controls of same stiffness group, black asterisks to 1kPa of same treatment group.

VSMCs are known to assemble podosomes in response to chemical or biophysical stimuli. Importantly, VSMC podosomes have been also identified in vivo and are a hallmark of phenotypic switching (*27*, *50*, *51*). Podosomes consist of a core of densely polymerized F-actin, surrounded by a ring complex, containing adhesion proteins. Indeed, using superresolution spinning disc microscopy we found cortactin, one of the podosome markers, localized with the F-actin core in VSMCs, which was surrounded by adhesion proteins zyxin or vinculin (Fig 1G,H). Localized degradations of the gelatin matrix were also observed around the podosome (Fig 1J,K). Surprisingly, the number of podosome-forming cells showed not only a strong increase after pressure stimulation compared to the control, but also depended largely on matrix stiffness, with a higher number of podosome forming cells on compliant surfaces. To confirm this unexpected result, we examined the effect of pressure and stiffness on primary bovine and human cells, which all recapitulated the observed changes (Supplementary Figure 1). Noticeable was a blunted effect on one human cell isolate due to a higher podosome forming activity under control conditions. Since these cells were already cultured for >10 passages, this was likely due to onsetting dedifferentiation and senescence. In terms of matrix remodelling, there were more podosome-forming (A7r5) cells and greater gelatin degradation on the 1kPa surface, compared to 130kPa. (Fig 1I and K). The extensive changes to the cytoskeletal structures were accompanied by a reduction in the cellular stiffness (mean±SEM: 3.78±0.24 kPa vs 2.21±0.18 kPa in control and HT treated cells) which was measured by nanoindentation with a large-diameter spherical tip placed approximately above the nucleus (Fig 1L, tip radius: 50μm, k=0.5N/m).

To further investigate the different behaviour of VSMCs we performed quantitative proteomics analysis on soft (1kPa) or stiff (130kPa) surfaces in presence or absence of cyclic hypertensive pressure. Abundance of smooth muscle proteins (including smooth muscle actin, metavinculin, caldesmon 1 or calponin) confirmed retention of a smooth muscle phenotype of the A7r5 cells (Suppl. Table 1). Hierarchical analysis of 1026 quantified proteins indicated primarily clustering of pressure treated samples vs control samples (Figure 2A). Individual pairwise analysis identified no, or only few significant changes between 1kPa control vs 130kpPa control, 1kPa HT vs 130kPa HT, or 130kPa control vs 130kPa HT treated samples. In contrast the pairwise comparison between 1kPa control and 1kPa HT pressure treated sample indicated 123 proteins that were significantly up or down regulated (FC>2, p<0.05) (Fig 2B-E and supplementary tables 2-5). The differential regulated proteins included 14 podosome associated proteins, 9 other cytoskeletal proteins, 10 stress response proteins (intriguingly all down regulated with pressure stimulation) and importantly 17 proteins that were previously associated with atherosclerosis or neointima formation (Figure 2F-K and supplementary note 1).

**Figure 2:**
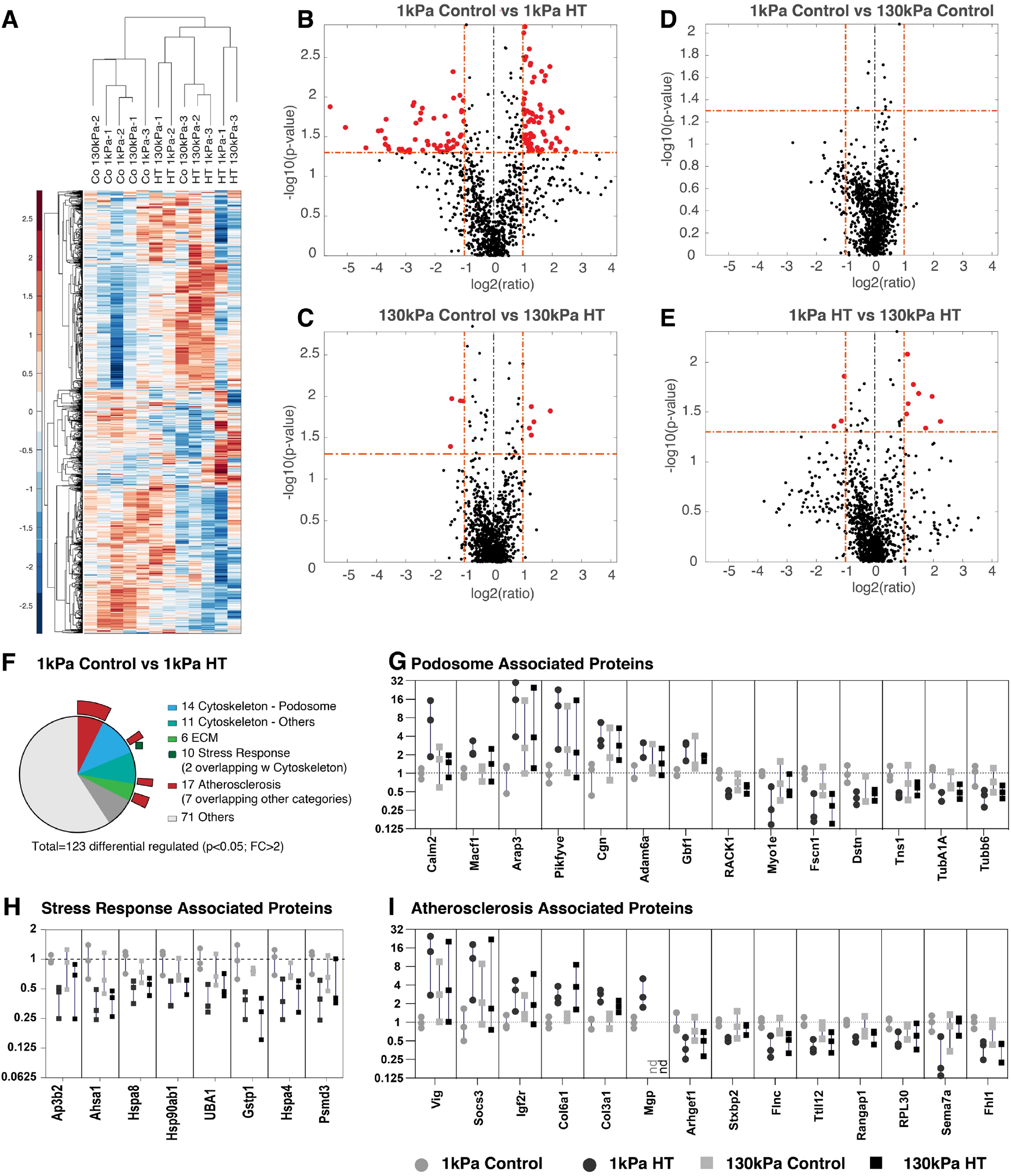
Quantitative proteomics analysis indicates stiffness and pressure dependent changes. A) hierarchical clustering of identified proteins indicates primary clustering of HT treated samples and control samples. Pairwise differential regulation analysis indicates 123 significant changes (FC>2, p<0.05; shown in red color) between 1kPa control and HT treated samples (B), compared to no, or only few significant changes for the other comparisons. (C-E). F-I) Literature and GO analysis of the identified 123 differential regulated proteins identified 14 proteins associated with podosome formation (G), 11 other cytoskeletal proteins, 6 extracellular matrix components, 10 proteins associated with stress response (H) and 17 proteins associated with Atherosclerosis (I), see also supplementary note 1; numbers including overlaps, see F, outer circle; G-I: alpha and beta tubulin as well as Myo1e are shown in G, but are also associated with atherosclerosis; alpha and beta tubulin also with stress response. From left to right: grey dots: 1kPa control; black dots: 1kPa HT; grey squares: 130kPa control; black squares: 130kPa HT;

Together these results suggest that the HT pressure globally alters the cell morphology and the actin organisation, but cells on the softer surface mimicking the stiffness of (pre-) atherosclerotic lesions show a distinct increase in the formation of matrix-degrading podosomes.

### Substrate stretching is insufficient to induce the podosome formation

The haemodynamic pressure results in compression and stretching of the extracellular matrix and cells in the arterial wall. The distension along the inner circumference of most human arteries was measured between 5 and 10% (*36*, *52*, *53*), whereby distension decreases in hypertensive patients, because of the accompanying arterial macro-scale stiffening (*42*). Therefore, we applied 10% cyclic biaxial stretch regime at the same frequency and duration of the pressure experiment (0.5Hz, 12h) to test if stretch alone is sufficient to induce the observed changes. After analysing the VSMCs with the same image analysis pipeline (see methods), we found however no clear differences between the cell populations (Supplementary Figure S2C,D). While we detected small but significant changes of cell shape (Supplementary Figure S2E-H), including an increase in eccentricity (more elongated shape) and form factor (more irregular shape), as well as a reduction in actin alignment, we did not detect changes in the cells ability to form podosomes (Supplementary Figure S2I). Similarly, 30 minutes static pressure application (180mmHg) stimulated podosome formation with a strong stiffness dependence, but we detected no change in the ability to form podosomes after 30 minutes static biaxial stretch (5 or 10%, Supplementary Figure S3).

### Vascular smooth muscle podosomes are mechanosensitive after induction through chemical stimuli

In addition to hypertensive blood pressure, disturbed flow and EC-VSMC crosstalk, through various PKC-activating signals from the ECs, are powerful stimuli for intima lesion formation and atherogenesis(*11*, *54–58*). Phorbol 12,13-dibutyrate (PDBu), a potent activator of PKC signalling in smooth muscle(*59*, *60*), has been previously used to induce the formation of podosomes in VSMCs(*61*, *62*). Consistently, we found a strong induction of podosome formation after PDBu treatment (Fig 3A,B). In particular, we detected a significantly higher fraction of podosome-forming cells, as well as significant differences in the turnover and function of podosomes on the substrate with lower stiffness (Fig 3C-E). Over time, podosomes were forming and turning over only once, or frequently cyclically appearing and disappearing at the same location (fluctuating podosomes, Fig 3C,D & Supplementary Movies 1&2). Moreover, podosomes often appeared in groups whereby the formation and disappearance followed a wave-like pattern, suggesting the cross talk and time delayed coordination among the podosomes. While we did not detect an obvious correlation between such wave pattern and stiffness, we found that podosomes on the soft surfaces turned over more rapidly (Fig 3E) and importantly, were degrading the extracellular matrix more efficiently on soft surfaces compared to cells plated on higher stiffness (Figure 3F,G). Together these results suggest that PDBu treatment leads to highly efficient activation of podosome formation, but the stiffness-dependent behaviour persists.

**Figure 3:**
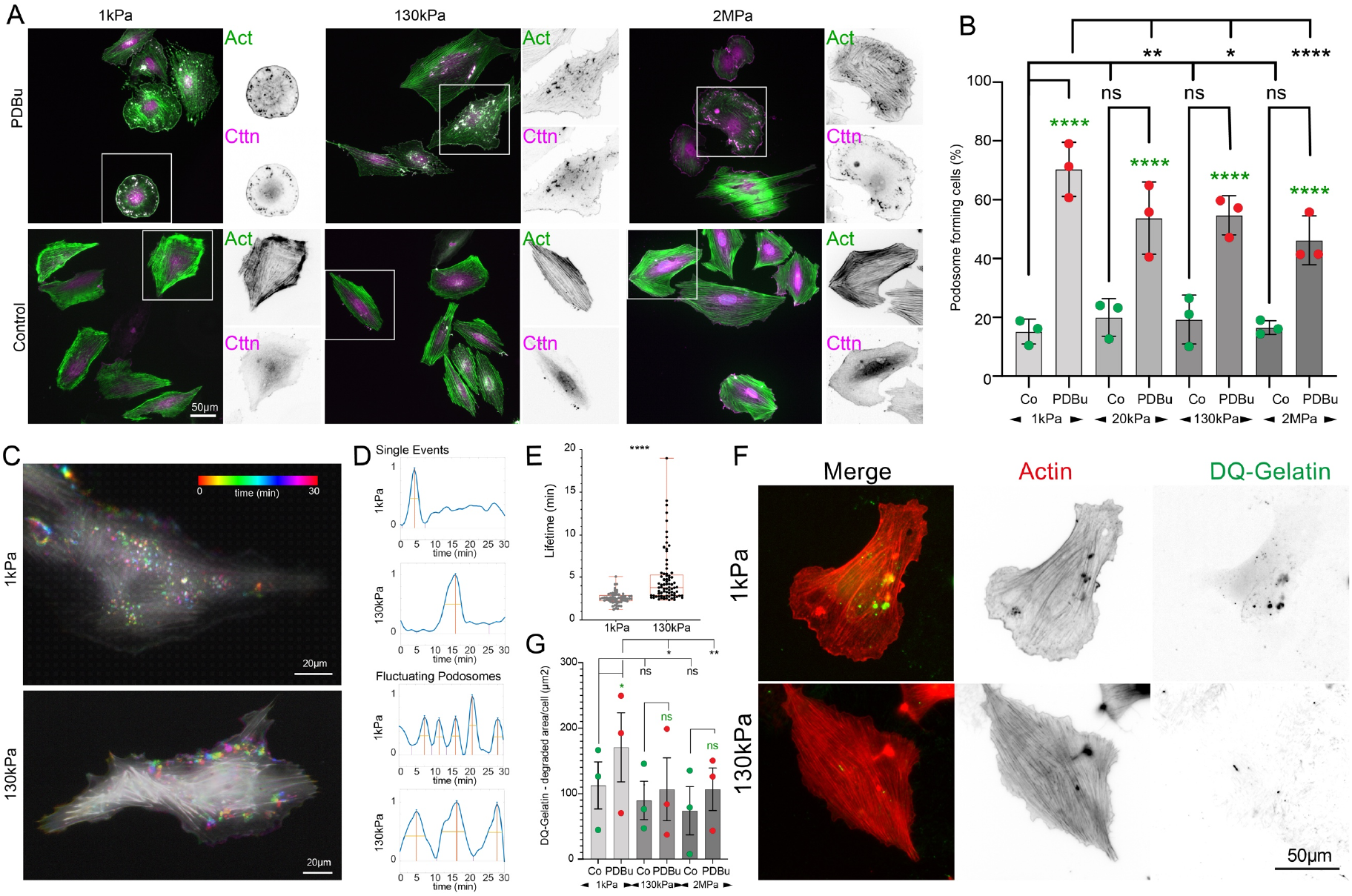
PDBu treatment leads to efficient podosome formation on all stiffnesses but podosome behaviour and matrix degradation are controlled by stiffness. G,H) PDBu treatment leads to stiffness dependent podosome formation (Fraction of podosome-forming cells, displayed as average per repeat, from three independent experiments with 10 images each and ~10-20 cells per image). I-J) Podosomes lifetime depends on substrate stiffness. I) Colour-coded time projection of VSMC treated with PDBu on 1 and 130kPa shows dynamic podosome formation (see also Supplementary Movies 1&2). J) Podosome formation (independent of stiffness) was detected as either single event or cyclically fluctuating podosomes but showed stiffness dependence in their turnover time (measured at half-maximum of the peaks) with an increased lifetime on higher stiffness (K). L,M) Gelatin degradation was more extensive on compliant surfaces (DQ-gelatin positive area per cell, displayed as average per repeat, from three independent experiments with n=30-100 cells analysed per repeat and condition). *p < 0.0332; **p < 0.0021; ***p < 0.0002; ****p < 0.0001; ns, not significant; p-values from ANOVA and Tukey test for multiple comparisons or unpaired t-test (E); H) green asterisks represent comparison to the controls of same stiffness group, black asterisks to 1kPa of same treatment group.

### Podosomal actin turnover in VSMCs is controlled by substrate stiffness

Faster turnover and recurrence of podosomes on compliant substrates suggested a stiffness dependent F-actin assembly. F-actin assembly at the podosome core is regulated through cortactin (via WASP and Arp2/3), or alternatively the actin-severing protein actin depolymerizing factor (ADF)/cofilin, which accelerates actin dynamics by generating additional barbed ends and has reported actin filament nucleation activity at high concentrations(*63*, *64*) The activities of cortactin and ADF/cofilin are regulated by the phosphorylation at Y421 and S3, respectively. We performed the western blotting with phospho-specific antibodies and found that levels of phosphorylated Y421 cortactin did not change with stiffness. However, there was a significant decrease in cofilin S3 phosphorylation on lower stiffnesses, suggesting a higher level of cofilin activity that was consistent with increased podosome dynamics (Fig 4A-C). Additionally, a stronger enrichment of cofilin at the podosome core was observed on softer surfaces (Fig 4D,E). To confirm the relevance of cofilin phosphorylation, we co-transfected mEOS2-actin and phosphorylation mutants of cofilin and measured the impact on the turnover of podosomal actin. After spot photo-activation, the dynamics of photo-converted mEOS2-actin at the podosome were monitored over time. Here, the presence of an active, non-phosphorylatable, cofilin mutant (S3A) increased actin turnover, while the presence of a phospho-mimicking S3E cofilin strongly reduced the turnover (Fig 4F-I). Together, these data suggest that cofilin is instrumental in regulating podosomal actin turnover downstream of extracellular matrix stiffness sensing.

**Figure 4:**
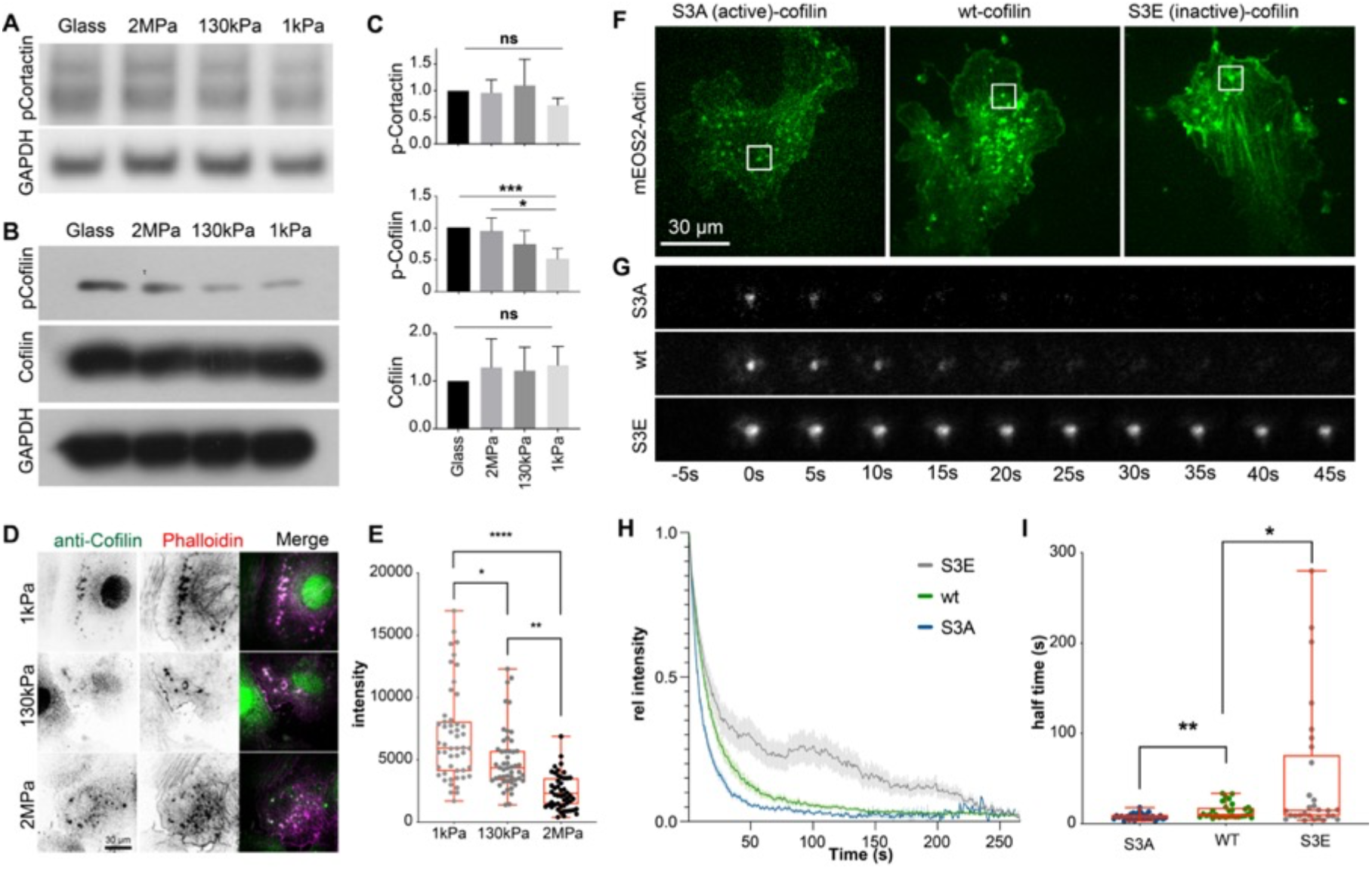
Cofilin localisation and activity respond to stiffness and regulate podosomal actin turnover. AC) Cortactin phosphorylation is unaffected by stiffness, but cofilin phosphorylation is reduced on soft surfaces, indicating higher activity. Error bars: SD; Test for linear trend between stiffness and pCofilin levels: p=0.0001; D,E) Cofilin is enriched at podosomes on soft surfaces. F-I) Transfection of phosphomimickinig or non-phosphorylatable cofilin regulates the turnover of podosomal actin. F) Cells were transfected with mEOS2-Actin and iRFP-cofilin wt, S3A (active) or S3E (inactive). G) Actin turnover was measured after photoswitching of mEOS2-Actin, indicating changes in lifetime. F-I) Fluorescence loss after photoconversion indicates slower podosomal actin turnover in presence of S3E and faster turnover in presence of S3A cofilin mutants. Plot of mean and SEM of 30 podosomes from 10 cells per condition (H) and half time (I). *p < 0.0332; **p < 0.0021; ***p < 0.0002; ****p < 0.0001; ns, not significant; p-values from one-way ANOVA and Tukey test; Box plots are displayed as median, upper and lower quartile (box) and range (whiskers).

### Podosomal cofilin is regulated through RhoA, ROCK2 and LIMK2 in a stiffness-dependent manner

LIM domain kinases (LIMKs) are known to directly phosphorylate cofilin at S3 and to reduce its actin severing activity(*65*). The activation of LIMK can be positively regulated by ROCK or PAK mediated phosphorylation. We sought to investigate how cofilin phosphorylation is regulated downstream of mechanical stimuli. In PDBu-stimulated VSMCs, we found that the level of phosphorylated LIMK showed the same trend as that of phosphorylated cofilin, and higher levels of phosphorylation were observed on the stiff surface (Fig 5A). Since the phospho-antibody detects both LIMK1 and LIMK2, we knocked down each isoform and found that LIMK2 was responsible for phosphorylating cofilin in A7r5 VSMCs (Fig 5B, Supplementary Figure S4 for knockdown validation). Next, we investigated the potential upstream kinases to promote the LIMK phosphorylation. We found that the activities of group 1 PAK (PAK1/2/3) and group 2 PAK (PAK4/5/6), detected by the respective phosphospecific antibodies, did not demonstrate any stiffness dependence. However, when ROCK1 and ROCK2 were knocked down individually, we found a strong reduction in cofilin phosphorylation after the knockdown of ROCK2 (Fig 5A,B and Supplementary Figure S4 for knockdown validation). In agreement with the effect of the knockdowns, pan-inhibitions of Rho kinases with Y27632 (20μM) and H1152 (10μM) or ROCK2-specific inhibition with KD025 (10μM) resulted in the decrease of both cofilin and LIMK phosphorylation (Supplementary Figure S5). In addition, we found that both LIMK2 and ROCK2 were colocalised and specifically enriched at the VSMC podosome (Supplementary Figure S6). When ROCK2 and LIMK2 were individually knocked down, the transwell migration of PBDu-treated VSMCs was strongly reduced (Fig 5C). Reintroductions of ROCK2 and LIMK2 in the respective knockdown condition restored the migration capability. RhoA is known to act as an upstream messenger to activate ROCK-mediated signalling transductions. Indeed, we further identified higher proportions of GTP-bound RhoA when VMSCs were plated on the stiff substrate (Fig 5D). Intriguingly, using a RhoA FRET activity sensor we found high RhoA activity in areas surrounding the podosome core (Fig 5E-H, Supplementary Figure S7 and Supplementary Movie 3). Initially, the RhoA activity was reduced during the podosome assembly, as predicted previously (*66*, *67*). However, flashes of RhoA activity increasingly and consistently appeared in the vicinity of the podosome core during the podosome disassembly (Fig 5F-I).

**Figure 5:**
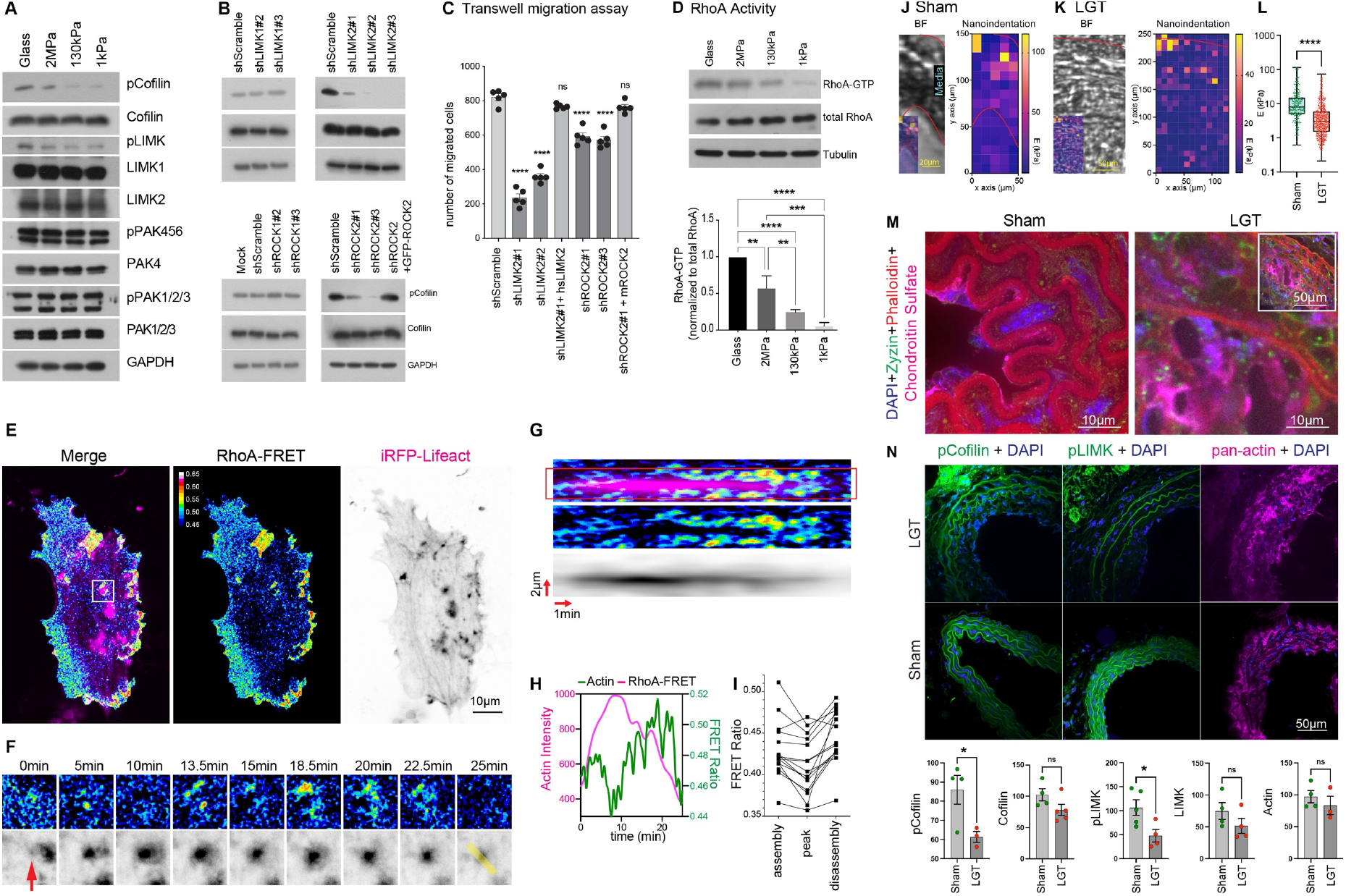
Rigidity sensing controls Cofilin activity through RhoA-ROCK-LIMK signalling. A) Western blot analysis of A7r5 VSMCs seeded on different stiffnesses. p-Cofilin and p-LIMK are reduced on compliant surfaces, while other antibodies do not detect stiffness dependent changes. B) LIMK2 and ROCK2 knockdown, but not LIMK1 or ROCK1 knockdown reduce p-Cofilin levels in A7r5 VSMCs (see Supplementary Fig S3 for knock down validation). C) LIMK2 and ROCK2 knockdown reduce transwell migration of VSMCs. Error bars: SEM; D) RhoA-GTP pull down assay shows a reduction in active RhoA on compliant surfaces (quantification from three independent repeats; Error bars: SD; Test for linear trend between stiffness and RhoA-GTP levels: p<0.0001). E-G) A7r5 cells were transfected with a RhoA-FRET biosensor and iRFP lifeact to analyze the dynamic changes to RhoA activity after PDBu stimulation. CFP/YFP images were taken every 5s after donor excitation with a 445nm laser and with a dual camera setup for simultaneous acquisition (see also Supplementary Fig S7). iRFP images were taken every 100s as a reference for the podosome position. F) Time points of area outlined in white in E. G) Kymograph over line marked in yellow in F. H) Intensity profiles over time of a 3μm wide line marked by outlines in G. I) FRET intensity at half-max actin intensity during assembly and disassembly as well as at peak intensity from n=15 podosomes from 3 independent experiments. The iRFP image stack was filtered with a 3D gaussian filter (x=1, y=1, z=10) for visualization purpose in panel G,H. J-L) Nanoindentation measurements indicate a reduced elastic modulus during neointima formation. J,K) BF images before nanoindentation (inset: overlay with measured Young’s modulus) for sham treated animals (J) or after carotid artery ligation (K: LGT); Quantified in (L); M) The increased compliance coincides with increased proteoglycan expression (chondroitin sulfate staining). Inset for LGT shows larger area for overview. N) frozen sections from sham treated mice and animals that underwent carotid artery ligation, stained for p-cofilin, pLIMK and pan-Actin together with DAPI. Quantification of the staining intensities of contractile VSMC in sham and ligation (LGT) samples indicate a reduction of both p-cofilin and pLIMK, while actin, cofilin and LIMK intensities are unaffected. Only cell areas were included in the quantification, which was not affected by typical background staining from the elastin layers. Data presented as mean per animal from n= 221 and 163 cells (Sham and LGT respectively, pCofilin); 274 and 330 cells (Cofilin); 288 and 142 cells (pLIMK); 195 and 238 cells (LIMK); 311 and 246 cells (Actin); *p < 0.0332; **p < 0.0021; ***p < 0.0002; ****p < 0.0001; ns, not significant; p-values from ANOVA and Tukey test for multiple comparisons (C,D) or unpaired t-test (L,N).

Apart from RhoA, Cdc42 is also reported to promote cofilin phosphorylation through PAK-mediated LIMK phosphorylation(*68*). VSMCs with Cdc42-GTP inhibition by ML141 (10μM) indeed exhibited a lower level of cofilin phosphorylation but did not show the deceased phosphorylation of PAK and LIMK, suggesting a non-canonical effect of Cdc42 inhibition on cofilin (Supplementary Figure S5). Together, these results show stiffness dependent and dynamic changes to RhoA activity which regulate ROCK2 and LIMK2 mediated cofilin phosphorylation.

Next, we sought to examine the link between matrix rigidity, LIMK and Cofilin phosphorylation in vivo. Previous studies, examining lipid rich regions in atherosclerotic plaques indicated that the intima might also undergo softening during its remodelling and neo-intima formation(*12*, *18*, *29*, *32*). Mouse carotid artery ligation is a well-established model for intimal lesion formation, whereby disturbed flow pattern and endothelial mechanosensing lead to neointima formation. We first assessed the mechanical properties of sham treated and carotid ligation tissues (LGT) by nanoindentation. Since the nanoindentation setup (500nm indentations with a probe with radius=50μm and k=0.5 N/m) resulted in a contact diameter of ~10μm, we spaced the individual measurements 10μm in radial direction and axial direction. Subsequently, we aligned the measurements with the bright field image to assign the stiffness map with the arterial wall area of the tissue sections (Fig 5I,J).

Our measurements (total of n=220 and 433 indentations from 3 animals each for sham and LGT), indicated in both cases a stiffness gradient from the luminal side to the outside of the artery. Overall, the measurements confirmed a reduced elastic modulus in the LGT model (mean ± SEM: 14.6 ± 1.3kPa vs 6.0 ± 0.4kPa; Figure 5I-K), coinciding with an increase in proteoglycan content as indicated by chondroitin sulfate staining (Figure 5L). While the nature of the samples did not allow us to identify the presence of podosomes, we nevertheless detected an accumulation of actin puncta and surrounding adhesion (zyxin) structures (Fig 5L, Supplementary Fig S8), consistent with previous reports of podosome formation in VSMCs in vivo (*50*). Importantly, we also detected a reduction in phospho-cofilin and phospho-LIMK staining in VSCMs in the neo-intima, while the actin, cofilin and LIMK staining intensities remained unaffected (Figure 5M).

Together, our results show stiffness dependent and dynamic changes to RhoA activity which regulate ROCK2 and LIMK2 mediated cofilin phosphorylation in vitro, and a correlation between stiffness and LIMK and cofilin phosphorylation in VSMCs in vivo.

### Pressure and PDBu stimulate cofilin dephosphorylation and podosome formation through Ca2+ and SSH

While the RhoA-ROCK-LIMK pathway regulated the stiffness dependence of cofilin activity and podosome formation, we also found that the level of cofilin phosphorylation was decreased after cyclic HT pressure stimulation (Fig 6A). However, the phosphorylation of LIMK remained unchanged, suggesting an alternative pressure-dependent mechanism to regulate cofilin phosphorylation. Mechanosensitive channels have received wide attention as regulators of cellular behaviour since their discovery in the 1980s and especially the cardiovascular system is highly dependent on the regulation through mechanosensitive ion fluxes(*10*, *69*, *70*). In particular, elevated Ca^2+^ levels have been shown to promote the phosphatase activity of slingshot, which dephosphorylates cofilin(*71*). Besides, PKC (activated through PDBu treatment) is not only modulated by Ca^2+^ but also affects Ca^2+^ handling(*72*). Therefore, we hypothesized that hydrodynamic pressure could lead to an increase in baseline Ca^2+^ levels to induce podosome formation. When VSMCs were subject to a single indentation with a maximum pressure of 24 ± 4.4kPa (180 ± 33mmHg, Fig 5B-E), we found a consistent increase in intracellular Ca^2+^ levels, reported by the calcium indicator Cal520. Similar to pressure stimulation, PDBu treatment also resulted in a consistent increase in intracellular Ca^2+^ levels (Fig 6F,G). Noteworthy, both the pressure and the PDBu stimulation only led to a slow increase, compared to the treatment with thapsigargin (TG), suggesting that both were inducing the influx of extracellular Ca^2+^ rather than the release from the intracellular calcium stores (Fig 6F,G). To examine the impact of Ca^2+^ influx on the cofilin phosphorylation, we found that the treatment with the divalent cation ionophore A23187 resulted in the decrease of cofilin phosphorylation, similar to PBDu (Fig 6H). When the phosphatase activity of slingshot was inhibited by Sennoside A(*73*), the level of cofilin phosphorylation increased (Figure 6I). Also, the effect of A23187 on phospho-cofilin was counteracted by simultaneous treatment with Sennoside A, whereas phosphorylation of LIMK was unaffected by SSH inhibition, either with or without A23187 (Fig 6I). Consistently, podosome formation (after HT pressure stimulation) was blocked after slingshot inhibition (either alone or in presence of Y27632), while VSMC efficiently formed podosomes on all stiffnesses after ROCK inhibition with Y27632 (Fig 6J). Together these results suggest that hydrodynamic pressure and PDBu treatment are both acting on cofilin phosphorylation through calcium and slingshot phosphatase to induce podosome formation, while stiffness sensing is modulating cofilin phosphorylation through RhoA-ROCK and LIMK activity (Figure 7). Since after SSH inhibition VSMCs lose the link between rigidity sensing and podosome formation (likely due to loss of intrinsic podosomal rigidity sensing - see also Figure Fig 3C-E and Fig 5E-H) we conclude that both signals are acting together to lead to a change in VSMC phenotype, podosome formation and extracellular matrix remodelling and ultimately will contribute to the disease progression.

**Figure 6:**
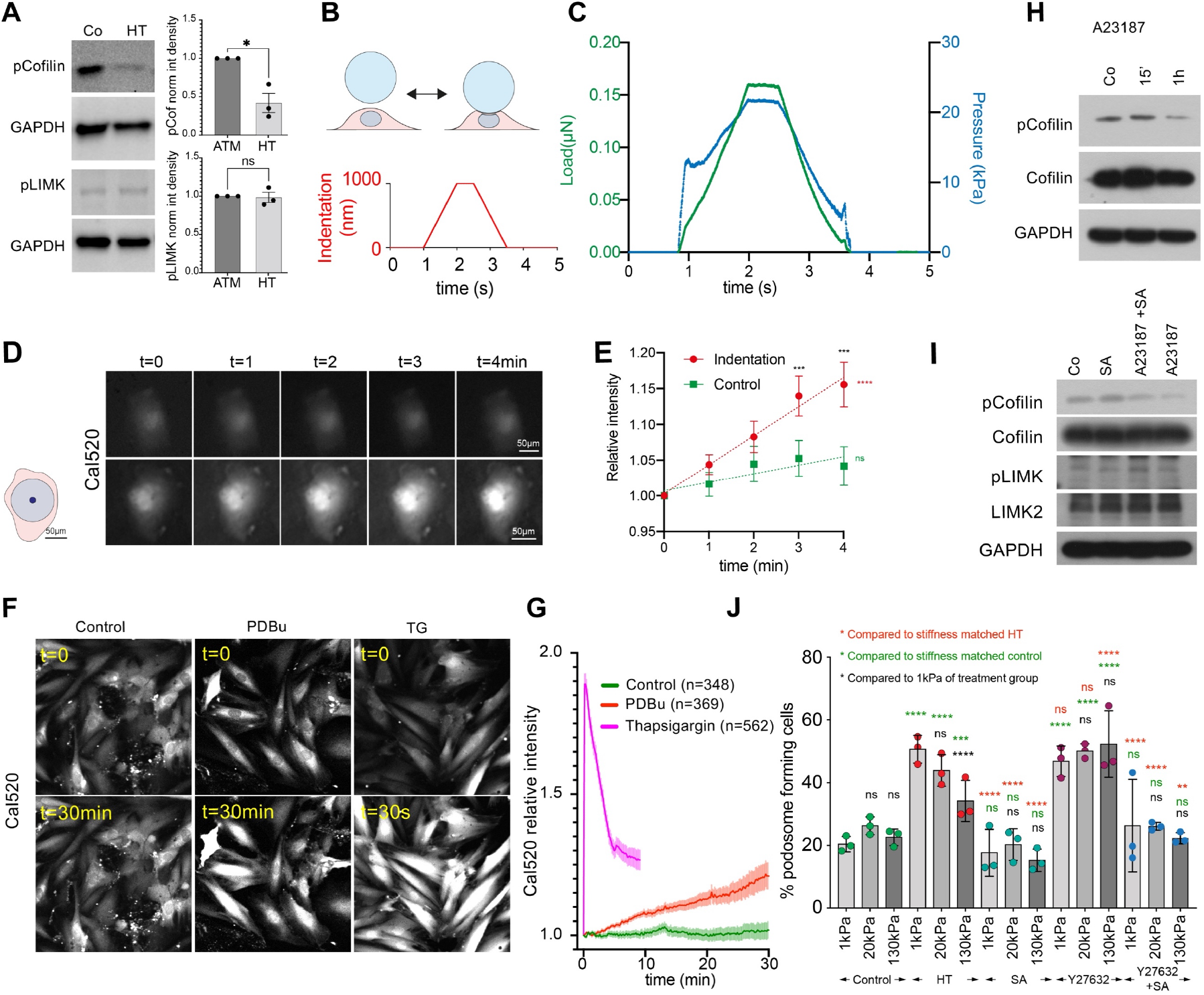
Pressure and PDBu stimulation induce calcium influx to regulate slingshot dependent cofilin phosphorylation. A) Hydrodynamic pressure stimulation reduces cofilin, but not LIMK phosphorylation. B) Schematic of pressure application with the nanoindenter (top) and indentation settings (bottom). A large diameter spherical tip is used to indent the cell with controlled pressure. C) example traces of corresponding load and pressure. Note that the initial large increase in load and pressure is due to the threshold load to find the cell surface. D,E) Indentation leads to a consistent increase in Cal520 intensity, while control cells do not show changes. Schematic on left indicates size of bead (in blue) and contact area at 1μm indentation (in dark blue) in comparison to cell area (in pink). E) Quantification of n=15 indented and 10 control cells (located nearby the indented cells) from 3 independent experiments. Error bars: SD; F,G) PDBu treatment leads to a consistent slow increase in Cal520 intensity, while thapsigargin (TG) induces a fast release of calcium from intracellular calcium stores. Note: lower panel shows 30 minute time point for Control and PDBu, but 30 second time point for TG; G) Quantification of n=348, 369 and 562 cells for control, PDBu and TG treatment. H) A23187 reduces cofilin phosphorylation after 1h. I) SSH inhibition increases cofilin phosphorylation after PDBu treatment and the reduction in pCofilin after A23187 is counteracted by simultaneous SSH treatment. J) SSH inhibition (either alone or together with Y27632) blocks podosome formation after hydrodynamic pressure stimulation on all stiffnesses, while ROCK inhibition enhances podosome formation after hydrodynamic pressure stimulation on all stiffnesses to a level otherwise seen on compliant surfaces only. *p < 0.0332; **p < 0.0021; ***p < 0.0002; ****p < 0.0001; ns, not significant; p-values from ANOVA and Tukey test for multiple comparisons or t-test (E); E) Black asterisks represent comparison for single timepoints and different conditions, green and red asterisks represent p-values from linear regression and test for deviation from zero. J: green asterisks represent comparison to the controls of same stiffness group, green asterisks represent comparison to the HT condition of same stiffness, black asterisks to 1kPa of same treatment group.

**Figure 7:**
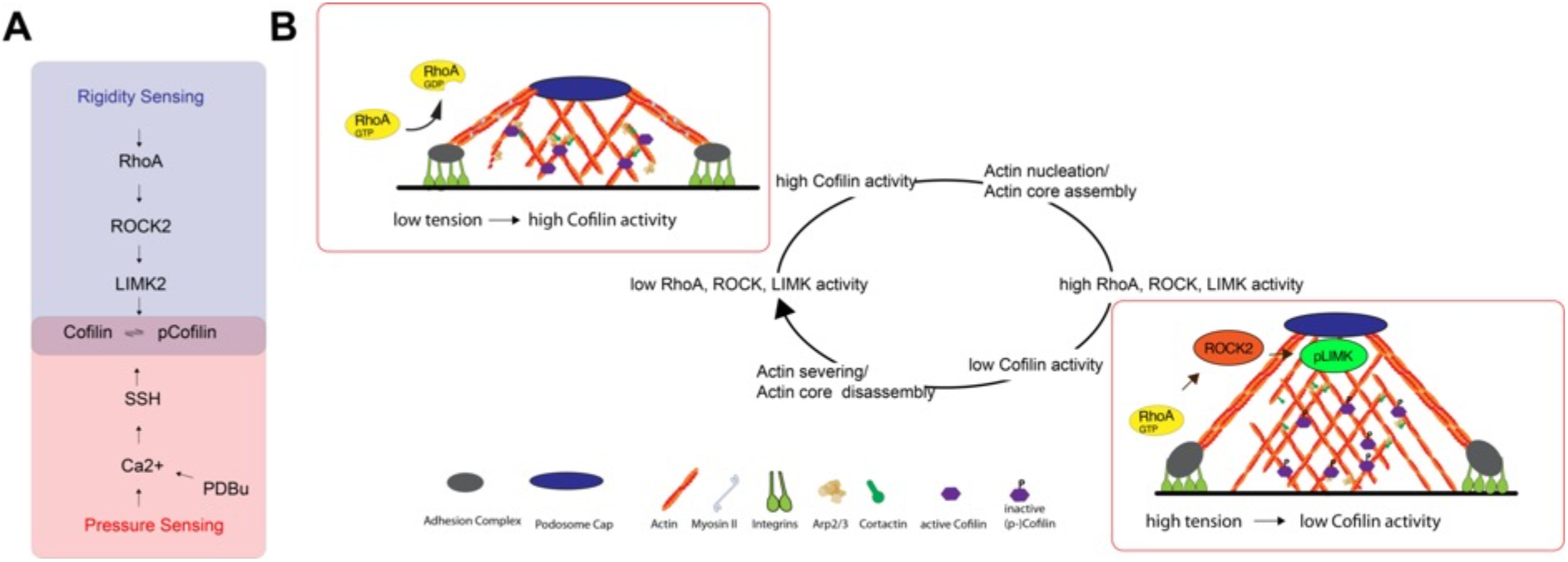
Schematic summarizing the pressure and stiffness dependent formation of podosomes in VSMCs. A) Pressure and PDBu signal through Ca2+ and SSH, while rigidity sensing acts through RhoA, ROCK2 an LIMK2 to modulate cofilin phosphorylation. B) Because cofilin switches between nucleation and severing depending on the concentration(*63*, *64*), fluctuations in cofilin activity lead to a cyclic nature of podosome formation: high cofilin activity will lead to actin nucleation and actin core assembly of podosomes, which eventually invokes RhoA activation at the podosome ring (through a currently unknown mechanism), thereby increasing ROCK and LIMK activity. This in turn reduces the cofilin activity, leading to actin severing and actin core disassembly and a reduction in RhoA activity.

## Discussion

Atherosclerosis is a severe cardiovascular disease. Mechanical stimuli have been strongly implicated in both disease onset and progression. The role of wall stress on endothelial cells is well documented and determines atheroprone and atheroprotective regions(*6–9*). Additionally, ageing (and associated changes to the mechanical properties of the arterial wall) and hypertension are major risk factors (*1–5*, *74*). Especially, the effect of blood pressure on the arterial wall changes during ageing and in disease. The enhanced macroscale stiffness leads to a reduced distension but increased radial stress and compression of the intima layer(*26*, *36*, *42*, *45*, *46*). These mechanical stimuli affect also the residing vascular smooth muscle cells (VSMCs), which are critically involved in the early stages of atherosclerosis (*12*). However, the effect of the different overlapping physiological or pathological mechanical stimuli on intimal VSMCs are still unclear.

Here we sought to investigate the role of the individual mechanical stimuli through hydrodynamic pressure stimulation of cells on surfaces with defined stiffness, as well as separate cell stretching experiments. This way, we could show that two mechanical stimuli, when combined, compliance (mimicking the intima or pre-atherosclerotic lesions), and hypertensive pressure lead to a phenotypic switch of vascular smooth muscle cells. Importantly this effect can be observed for the A7r5 VSMC line, as well primary bovine and human vascular smooth muscle cells. In all cases, a partial phenotypic change can be detected independent from substrate stiffness. However, a complete phenotypic switching including maximal formation of podosomes and associated matrix degradation is strongly dependent on a combination of pressure treatment (or chemical stimulation) with a compliant environment. This was further confirmed by our proteomic data set, which only picked up significant different protein levels between pressure treated and control samples, when these were seeded on compliant surfaces. Consistent with the observed phenotypic change, we detected a range of molecules that were previously associated with podosome formation and/or atherosclerosis amongst the differential regulated proteins.

The implication of this is significant for the processes seen during early atherosclerosis. The increased expression of non-collagenous extracellular matrix in the vessel wall during the early atherosclerotic phase further modifies the microscale stiffness, also measured here in a neointima model. Hence, the combination of compliance with hypertensive pressure will enhance the phenotypic switch and accelerate matrix remodelling and atherosclerotic progression in a downward spiral.

It is important to mention that our findings are not contrasting the importance of endothelial mechanosensing in onset and progression of atherosclerosis (*6–9*, *11*). Rather, our data suggests that chemical signals sent from the endothelium can initiate the phenotypic switch in VSMCs through various chemical signalling pathways, including PDGF and Wnt pathways that activate PKC (*11*, *54*, *55*) and thus will affect calcium influx (*72*), cofilin phosphorylation (*71*) and podosome formation. Similarly, hypertension and atherosclerosis have been both linked to an upregulation of the vasoconstrictors Endothelin-1 (*56*) and Angiotensin II(*57*), which likewise lead to increased production of diacylglycerol and downstream activation of PKC (*58*) and indeed PKC is deregulated in atherosclerosis(*75*). This shows that mechanical and chemical signals are not mutually exclusive but rather act together to speed up the disease progression, whereby the downstream formation of podosomes contributes to the remodelling of the pre-atherosclerotic lesions. Mechanosensitive behaviour of podosomes was previously studied in monocytic cells, with differing outcomes. One study using dendritic cells on a fibrinogen matrix found an increase in podosome formation with increasing stiffness, while another study using macrophages found a higher degree of podosome formation on compliant fibronectin coated hydrogels (*76*, *77*). Similar to the latter study we find a larger degree of podosome formation and dynamicity on soft fibronectin, gelatin or collagen coated (not shown) surfaces. Noteworthy, we also detect a fluctuating appearance of podosomes at the same locations, whereby the oscillation period matches well with force oscillations that were previously observed in podosomal force studies (*76*, *78–80*).

A slow oscillating component was previously reported in the range of 7 ± 4 minutes, which was matching the actin assembly period at the podosome core (*80*, *81*). Fluctuating actin assembly in turn is consistent with the dual activity of cofilin as nucleator and severing protein, depending on its (active) concentration (*63*, *64*), regulated downstream of RhoA signalling. As the substrate stiffness orchestrates RhoA activity, the downstream pathway of ROCK and LIMK will affect cofilin phosphorylation and the overall propensity of cells to initiate podosome formation.

Indeed, the level of RhoA-GTP is decreased on compliant surfaces, leading to the cofilin mediated actin nucleation and a larger fraction of cells forming podosomes after pressure or chemical stimulation(*66*, *67*). Moreover, once cells start forming podosomes, the regulation of RhoA-GTP level can locally form a feedback loop to dynamically modulate the cofilin activity and myosin-dependent oscillation. The increase of RhoA-GTP level during the assembly phase of podosome can reduce the activity of cofilin in actin nucleation and lead to the subsequent podosome disassembly.

Together this suggests that stress or strain dependent effects might lead to the activation of RhoA and points towards a role for a negative regulator of RhoA activity in the podosome adhesion ring. At this moment we can only speculate about the upstream mechanosensor, but potential molecules include ARAP3, a dual GTPase activating protein (GAP) for RhoA and Arf, is dynamically recruited to the podosome in a FAK/PI3K-dependent manner (*67*) and our quantitative proteomics analysis found Arap3 strongly upregulated after pressure stimulation, especially on soft surfaces, consistent with a higher dynamicity of podosomes. Other BAR-domain containing RhoA GAPs, such as srGAP1, RICH1, and GMIP, might sense the change of membrane curvature during the podosome formation (which we indeed detect even on glass coverslips, where podosomes appeared to lift the membrane upwards in their vicinity, Supplementary Figure S9)(*82*) and regulate local RhoA activity(*83*, *84*). However, future experiments need to untangle the stiffness dependent regulation of RhoA activation at the adhesion ring.

Overall, our findings suggest a regulation of VSMC phenotypic switching through combined mechanical signalling through hemodynamic pressure and extracellular matrix stiffness, which could be targeted to mitigate early atherosclerotic processes and abate disease progression.

## Methods

### Antibodies and Reagents

Primary antibodies included vinculin (Sigma V9131, 1:200 for IF), cortactin (Sigma 4F11, 1: 100 for IF), phospho-cofilin (Ser3) (CST 3311, 1:3000 for WB), cofilin (D3F9, CST 5175, 1:5000 for WB, 1:100 for IF), phospho-cortactin (Tyr421)(ThermoFisher 44854G, 1:1000 for WB), phospho-LIMK1 (Thr508)/LIMK2 (Thr505) (CST 3841, 1:1000 for WB), LIMK1 (CST #3842 or BD Biosciences # clone 42, 1:1000 for WB), LIMK2 (Abcam ab45165 or CST #8C11, 1:1000 for WB, 1:100 for IF), PAK1/2/3 (CST 2604, 1:1000 for WB), phospho-PAK1/2/3 (Thr402) (ThermoFisher PA1-4636, 1:1000 for WB), phospho-PAK4 (Ser474)/PAK5 (Ser602)/PAK6 (Ser560) (CST 3241, 1:1000 for WB), PAK4 (CST 62690 or 3242, 1:1000 for WB), ROCK2 (D1B1, CST 9029, 1:1000 for WB), ROCK1 (C8F7, CST 4035, 1:1000 for WB), pan-actin (Merck #MAB1501), chondroitin sulphate (BD Biosciences #554275), β-tubulin (CST 2146, 1:3000 for WB), and GAPDH (ThermoFisher AM4300 or Abcam ab9485, 1:10000 for WB). Alexa Fluor 488 Phalloidin and Alexa Fluor 568 Phalloidin were from Invitrogen.

The following reagents were used at the specified concentrations after 24h serum starvation: Phorbol 12,13-dibutyrate (PDBu, Tocris #4153, 1μM), Thapsigargin (SCBT, sc-24017, 100nM), A23187 (SCBT, sc-3591, 1μM), SennosideA (Sigma 68909-5MG-F or Selleck #S4033, 10μM), Y27632 (Cambridge Bioscience or Selleck #S1049, 10 to 20μM), H1152 (Tocris #2414, 10μM), KD025 (Selleck #S7936, 10μM), ML141 (Selleck #S7686, 10μM), and A23187 (Selleck #S7778).

### Plasmids

Cofilin WT, cofilin S3A and cofilin S3E-pmiRFP were created by subcloning the respective cofilin constructs from pEGFP-N1 human cofilin plasmids using HindIII-HF and XmaI restriction sites.

pEGFP-N1 human cofilin WT, cofilin S3A and cofilin S3E were a gift from James Bamburg (Addgene plasmids # 50859, 50860 & 50861)(*85*). pmiRFP703-N1 and pLifeAct-miRFP703 were a gift from Vladislav Verkhusha (Addgene plasmid # 79988 & 79993)(*86*). mEos2-Actin-7 was a gift from Michael Davidson (Addgene plasmid # 57339)(*87*). GFP-ROCK2 was a gift from Alpha Yap (Addgene plasmid # 101296)(*88*). pLKO.1 - TRC cloning vector was a gift from David Root (Addgene plasmid # 10878)(*89*). Tractin-tomato and tractin-GFP(*90*) were a gift from from M. Schell. The RhoA-FRET biosensor was a gift from Oliver Pertz(*91*). mTFP-TRAF-Venus and mTFP-5AA-Venus were a gift from Nicolas Borghi(*92*). Vinculin-mTFP and Vinculin-Venus were a gift from Carsten Grashoff (*93*).

### shRNA

The following sequences, inserted into an empty pLKO.1-TRC vector, were used for shRNA knockdowns: shScramble: 5’-CCTAAGGTTAAGTCGCCCTCG-3’; shROCK1#1 5’-GGTTTATGCTATGAAGCTTCT-3’; shROCK1#2 5’-GCATTTGCCAATAGTCCTTGG-3’; shROCK1#3: 5’-GCCGACTTTGGTACTTGTATG-3’; shROCK1#4: 5’-GCACCAGTTGTG CCTGATTTA-3’; shROCK2#1: 5’-TGCAAAGTTTATTATGATATA-3’; shROCK2#2: 5’-AACGTGGAAAGCCTGCTGGAT-3’; shROCK2#3: 5’-GCAGAAAGTTCCAAACAGA-3’; shROCK2#4: 5’-GTAGAAACCTTCCCAATTC-3’; shLIMK1#1 “ 5’-CCTCCATTCGATGAACATCAT-3’; shLIMK1#2: 5’-AAGACTTGCGTAGCCTTAAGA-3’; shLIMK1#3: 5’-AATGCAGACCCTGACTATCTG-3’; shLIMK2#1: 5’-AATGGCAAGAGCTACGATGAG-3’; shLIMK2#2: 5’-AACAACCGAAATGCCATCCAC-3’; shLIMK2#3: 5’-GCCATCAAGGTGACTCACAAA-3’; all shRNA oligos were from Integrated DNA Technologies IDT. Cells were transfected with shRNAs for 96h before the experiments.

### Cell culture, Immunostaining and Microscopy

Primary VSMCs were obtained from explants of bovine aorta and from human aortic tissue from two healthy female donors aged 35 and 38 years as previously described(*94*). Human samples were obtained with written informed consent and approval from the Research Ethics Committee which conforms to the principles outlined in the Declaration of Helsinki.

A7r5 rat arterial vascular smooth muscle cells were cultured in DMEM with 5% FBS, 1% Glutamax and 1% Penecillin/Streptamycin (P/S)). Cells were transfected using Lipofectamine LTX with plus reagent, following the manufacturer’s instructions. For immunostaining, cells were fixed with 4% PFA for 10 minutes, permeabilized with 0.2% triton X-100 in PBS for 5 minutes, blocked with 5% BSA in PBS for 1h and stained in the antibody solutions in immunostaining buffer (20 mM Tris, 155 mM NaCl, 2 mM EGTA, 2 mM MgCl2, 1% BSA at pH 7.4) (*95*). Cells were washed three times for 10 minutes with PBS after each step and mounted in MOWIOL 4-88 (0.1g/ml in Glycerol/Water/Tris (0.2M, pH8.0) at a ratio of 1/1/2) containing a final concentration of 4% n-propyl gallate. Live cell imaging was performed on an inverted Nikon Eclipse Ti-E microscope with a Nikon DS-Qi2 sCMOS camera (Podosome lifetime imaging) or a Nikon Ti2 SoRa spinning disc microscope with two Photometrics Prime BSI cameras for simultaneous imaging (FRET, Calcium imaging, used in confocal mode), both equipped with environmental chamber with temperature and CO2 control. Podosome immunofluorescence imaging was performed on a or a Nikon Ti2 SoRa spinning disc microscope with a 60x 1.49NA objective in SoRa 4x magnification mode. Imaging of cells after pressure stimulation was done on a Leica DMI8 epifluorescence microscope with a Leica DFC9000 GT camera.

Fluorescence loss after photoconversion experiments were performed a Perkin Elmer UltraView VOX Spinning Disc Confocal. The photoconvertible mEos2-Actin was excited by an instant pulse of 405nm UV laser at cofilin-positive podosomes. The decay phase of the photoconverted red mEos2-Actin was then imaged at the rate of 1 second per frame.

### Animal model and tissue sections

Complete ligation of the left common carotid artery (LCCA) was performed with eight-week-old male C57BL/6 mice as described previously(*96*). After harvesting 28 days post-ligation, the proximal and distal 2-3 mm of LCCA were discarded and the remaining portion (4-5 mm) were embedded in optimal cutting temperature compound for frozen sectioning on a cryostat-microtome (Leica Biosystems). Frozen sections (10 μm thick) were prepared and subjected to immunofluorescence staining analyses with the indicated antibodies.

For this, frozen sections were thawed at room temperature for 15-20min and rehydrated twice with PBS for 5 min. Tissues were blocked for 30 min with 5% horse serum diluted in PBS and incubated with primary antibodies (1:100) at 4°C overnight. Next day, sections were washed with PBS for 5 min, three times and incubated with secondary antibodies (1:200) for 60 min at 37 °C. Tissues were washed three times with PBS, incubated for further 15min with DAPI (1:1000), washed again with PBS and mounted using Prolong Gold mounting media (ThermoFisher). Z-stacks of the 10μm thick sections were taken on a Nikon Ti2 SoRa spinning disc microscope with a 60x oil objective (unless otherwise specified). Maximum intensity projections were calculated and staining intensities of cells were measured using imageJ. Indicated number of cells (Figure 5N, figure legend) from 3 sections each from n=3-5 animals were included in the analysis and plotted as mean per animal.

### PDMS substrates

Flat PDMS substrates were prepared as described previously(*97*). Briefly, Sylgard 184, Sylard 527 or mixtures at the Ratios of 1:5, 1:10 and 1:20 were spin coated with a 150i spin processor (SPS), onto coverslips. Before spin coating, Sylgard 527 was pre-cured at 70°C for 30 minutes with intermitted mixing to achieve a comparable viscosity to the Sylgard 184 mixture. The stiffness of mixtures was measured by Rheology as described previously(*47*). PDMS substrates were coated with fibronectin, or DQ-gelatin for matrix degradation experiments.

### Pressure stimulation

Hydrodynamic pressure stimulation was performed in a MechanoCulture TR stimulator, placed inside a tissue culture incubator. The stimulator was modified setup for low pressure stimulation in the range of (human) normal (NBP) and hypertensive blood pressure (HT) and perfusion ports to supply pre-saturated cell culture medium. Cells were stimulated with a sinusoid profile (stretch: 0.5s, duration: 1s, hold: 0s, recovery 0.5s) and alternating between set pressures of 16 (peak load) and 8kPa (pre-load) to reach a measured pressure profile of 120/60mmHg for normal blood pressure stimulation and between 26 and 16kPa for a measured pressure profile of 180/120mmHg for hypertensive pressure stimulation. Cyclic pressure stimulations were performed for 12h and static pressure stimulations (at peak pressure setting) were performed for 30minutes. Control cells were placed inside the stimulator without applying pressure.

### Cell stretching

Cell stretching experiments were performed using a FlexCell-4000 setup. Cells were seeded onto BioFlex Collagen I plates at ~50% confluence for biaxial strain stimulation the following day. Cells were stretched using a sinusoid profile cycling between 0 and 5% or 0 and 10% strain. Unstretched control cells were seeded onto unstretched membranes in parallel, retained in TC incubator, and then fixed simultaneously once the experiment had finished.

### Nanoindentation

Nanoindentation experiments were performed using an Optics11 Chiaro nanoindenter attached to a Leica DMI-8 microscope. Cell measurements were performed above the nucleus with a R=50μm, k=0.5N/m probe (suitable for a stiffness range between approx. 0.5 and 80kPa). The Hertzian contact model was used to fit the data. The contact point was identified from a contact point fit of the data to 20% of the maximal load and used to subsequently fit the Young’s modulus using an indentation depth of 1μm.

For tissue sections, measurements were performed with a R=50μm, k=0.5N/m probe and 500nm indentations in the matrix scan mode. Images of the section with the probe in contact at the start and end position were taken for alignment of the measurements with bright field image. Only measurements overlapping with the arterial wall were included in the analysis.

### Ca2+ measurements

For Calcium measurements cells were plated on fibronectin coated PDMS coverslips and loaded with Cal520 AM (abcam, ab171868) according to the manufacturer’s instructions. Cells were then treated with 1μM PDBu, or 1μM thapsigargin and imaged on a Nikon Ti2 SoRa spinning disc microscope in confocal mode (50μm pinhole) with a Plan Fluor 10x air objective. Untreated cells were used as a control. To measure the calcium response after pressure stimulation, controlled force was applied onto cells using a Chiaro nanoindenter with a R=49μm, k=0.46N/m probe, using an indentation depth of 1μm (equalling a contact area of 150μm2) with a maximum pressure of 24 ± 4.4kPa.

### Gelatin degradation

DQ-Gelatin is a fluorogenic substrate which emits green light once degraded by matrix metalloproteinases and excited by 488λ. To build a thin layer of gelatin on PDMS, 25mm cover slips pre-coated with PDMS were treated with 18W air plasma for 1 minute and then immediately coated with 0.1% DQ-Gelatin solution (DQ-Gelatin From Pig Skin, Fluorescein Conjugate; D12054) at room temperature for 10 minutes in a light-protected sterilized hood. Excess gelatin solution was removed by PBS. A7r5 cells were seeded at ~50% confluence 1 day before the experiment (PDBu treatment of pressure stimulation). For quantification of DQ-Gelatin degradation, a Fourier bandpass filter was applied to increase the signal to noise ratio of the DQ-Gelatin channel. The filtered images were combined with the actin and DAPI channels and analysed using a cell profiler pipeline. Briefly, DQ-Gelatin positive areas were detected as primary objects, masked by the previously identified cell areas and then combined and measured as sum of the degraded area per cell.

### Image analysis

Image segmentation was performed using CellProfiler, ImageJ (OrientationJ) and Matlab as described previously(*98*). Additionally, we added actin segmentation to the CellProfiler pipeline and trained an Ilastik pixel classification to detect actin and cortactin dots. All measurements were combined, and dimensionality reduction was performed in Matlab using the t-SNE algorithm. Cluster borders were drawn manually, and populations from each separate repeat were calculated. Additional cell morphological analysis was performed with the visually-aided morpho-phenotyping image recognition “VAMPIRE” software(*99*). For analysis of Fluorescence loss after photoconversion, mEos2-Actin intensity decay profiles of each podosome were measured by ImageJ and fitted into a one-phase decay curve. FRET analysis was performed in ImageJ. First donor and acceptor bleed through were determined using Vinculin-mTFP and Vinculin-Venus constructs using the PixFRET plugin. Donor bleed through was best fit with a constant value and Acceptor bleed through was determined to be negligible. Movies were processed using imageJ macro, by subtracting the background from all channels, calculating the bleed through corrected donor image, subtracting the bleed through corrected donor image from the FRET image, and then normalizing this by the Donor intensity.

### RhoGTP pull down

A7r5 cells were seeded on the newly prepared fibronectin-coated PDMS substrate with the defined stiffness for 48 hours and were then treated with 1μM of PBDu for two hours. To detect the RhoA-GTP level, cells lysates were prepared according to manufacturer’s instructions (RhoA Pull-Down Activation Assay Biochem Kit; Cytoskeleton Inc., BK036). Specifically, lysates with 300μg protein were incubated with the Rhotekin-RBD containing agarose bead at 4°C on a rotator. Total RhoA in the cell lysate and activated form of Rho-GTP in the agarose bead were detected by Western blot.

### Quantitative Proteomic analysis

Was performed at the proteomics facility Denmark Hill, King’s College London. Samples were loaded onto 1D SDS gel for in-gel reduction, alkylation and digestion with trypsin, prior to subsequent analysis by mass spectrometry. Digested peptides were labelled with TMT16plex tags, according to the protocol provided by the manufacturer.

Each TMT set sample was resuspended in re-suspension buffer (2% acetonitrile in 0.05% formic acid) to be analysed by LC-MS/MS with triple injections. Chromatographic separation was performed using an Ultimate 3000 NanoLC system (Thermo Fisher Scientific, UK). Peptides were resolved by reversed phase chromatography on a 75μm*50cm C18 column using a three-step gradient of water in 0.1% formic acid (A) and 80% acetonitrile in 0.1% formic acid (B). The gradient was delivered to elute the peptides at a flow rate of 250nl/min over 100 min.

The eluate was ionised by electrospray ionisation using an Orbitrap Fusion Lumos (Thermo Fisher Scientific, UK) operating under Xcalibur v4.1. The instrument was programmed to acquire using a “Synchronous Precursor Selection with MultinotchMS” method (SPS) for accurate and sensitive quantitation based on isobaric TMT tags.

Raw mass spectrometry data were processed into peak list files using Proteome Discoverer (ThermoScientific; v2.5). Processed data was then searched using Sequest search engine embedded in PD 2.5, against the current version of the reviewed Swissprot *Rat* database downloaded from Uniprot.

The LC-MS/MS analysis successfully identified over 10,000 protein groups containing at least 1 peptide identified across all samples with a peptide cut-off threshold of FDR0.05. After peptide sequence identification, default thresholds (Co-Isolation threshold: 50; Average S/N threshold: 10; SPS Mass Matches threshold: 65) were applied during the reporter quantification to remove redundant peptide spectrum matches (PSMs). Proteins and peptides were then quantified based on reporter ion ratios (signal to noise) of non-redundant PSMs. Among over 10,000 identified protein groups, 1026 protein groups were quantified. Since the peptide cut-off was set up as FDR 0.05, a further filtering with a Xcorr score greater or equal to 1.5 and valid present over 70% across replicates within each condition was applied, aiming to remove poor identification and quantification. As a result, 1026 quantified protein groups are used for downstream differential analysis (Supplementary tables 1-5). Cluster analysis and pairwise differential analysis was carried out using the Matlab clustergram using the correlation distance metering and mavolcanoplot functions.

### Statistical analysis

Data sets were tested for normal distribution using the Shapiro–Wilk test. All statistic tests were performed with GraphPad Prism using either t-tests for two conditions, or ANOVA and correction for multiple comparisons.

## Author Contributions

PS, BS, ZF, EM, MW, IX, SZ, XZ and TI performed the experiments and analysed the data. GJ, CS, CHY and TI designed the study. QX, CHY and TI supervised the work and CHY and TI wrote the manuscript.

## Acknowledgments

We thank David Rumschitzki and Peter Weinberg for helpful discussions and input on arterial and intima compressibility. We would especially like to acknowledge M. Schell for the Tractin plasmids, Oliver Pertz for the RhoA biosensor plasmids, Nicolas Borghi for the mTFP-TRAF-Venus and mTFP-5AA-Venus, and Carsten Grashoff for the Vinculin-mTFP and Vinculin-Venus plasmids. We would like to thank Pei-Hsun Wu and Dennis Wirtz for the GUI/code and help with the VAMPIRE tool. We would like to thank BBSRC (BB/S001123/1) and BHF (PG/20/6/34835) for generous support.

