## Supplementary Information and Figures for "Matrix stiffness and blood pressure together regulate vascular smooth muscle cell phenotype switching"

### Supplementary Figures:

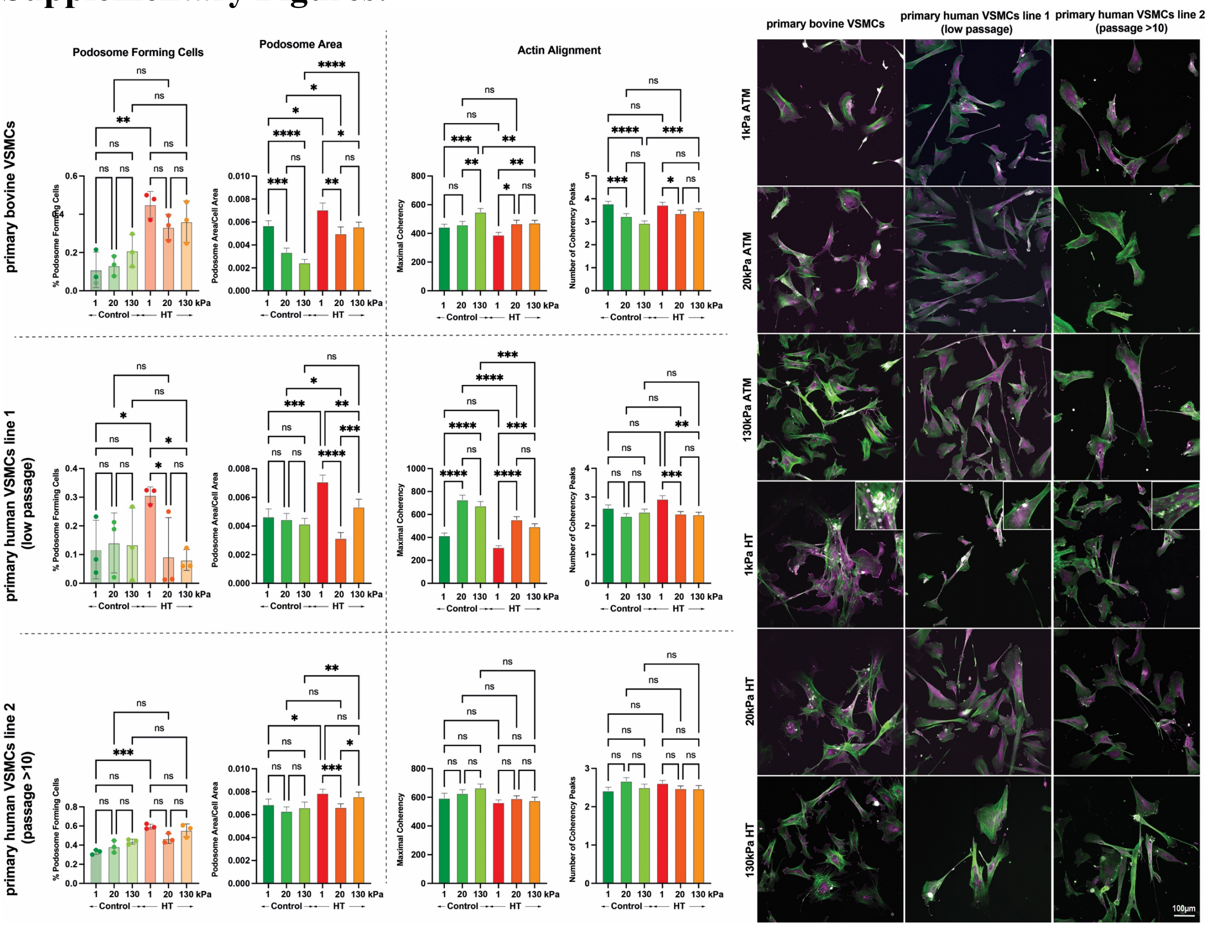


**Supplementary Figure S1:** Combined compliance and cyclic hypertensive pressure result in podosome formation and actin rearrangements also in primary bovine and human vascular smooth muscle cells. Cells were plated on PDMS coated coverslips with different stiffness (1,20,130kPa) and placed under cyclic hydrodynamic pressure mimicking hypertensive blood pressure (180/120mmHg). Cells were then fixed, stained. F) HT treatment resulted in reduced actin organisation on 1kPa (Coherency measurements using the orientationJ plugin), as well as podosome forming cells and podosome area (pooled data from three separate experiments). Insets at 1kPa HT display some examples of podosome forming cells.

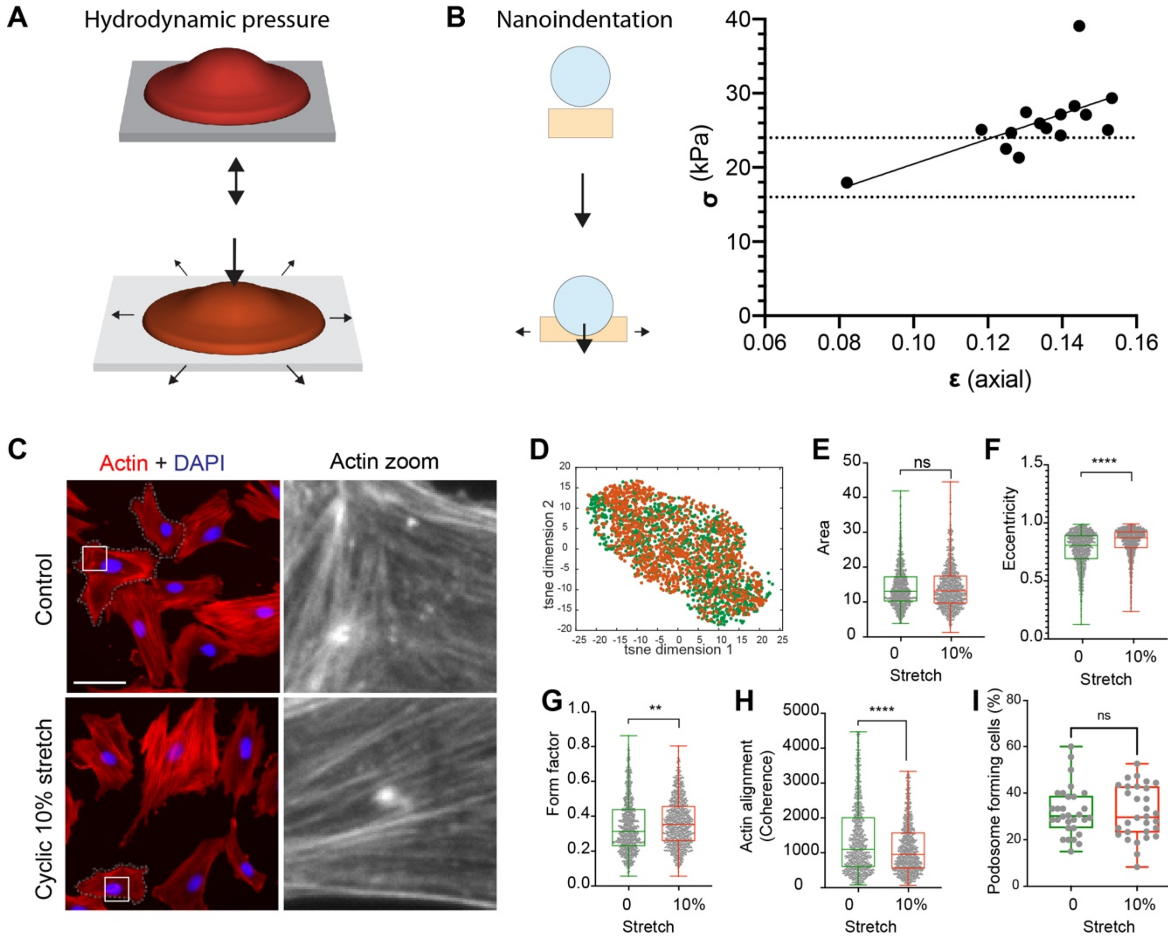


**Supplementary Figure S2: Cell Stretch alone is not sufficient to induce podosome formation.** A) schematic showing pressure stimulation will lead to compression and stretch of cells (red) and PDMS (grey). B) Nanoindentation was used to determine the amount of axial strain after applying stress in the range of hypertensive pressure. ~24kPa stress resulted in 12% axial strain of a 130kPa PDMS gel, equivalent to approximately 6% transverse strain. C-I) Cells were stained with Phalloidin and DAPI and analysed with the image analysis pipeline described in Figure 1. D) Both stretched and control cells clustered together. E-H) only small differences were observed for Eccentricity, Form Factor and Actin alignment. I) The fraction of podosome forming cells was unchanged after 12h cyclic stretch.

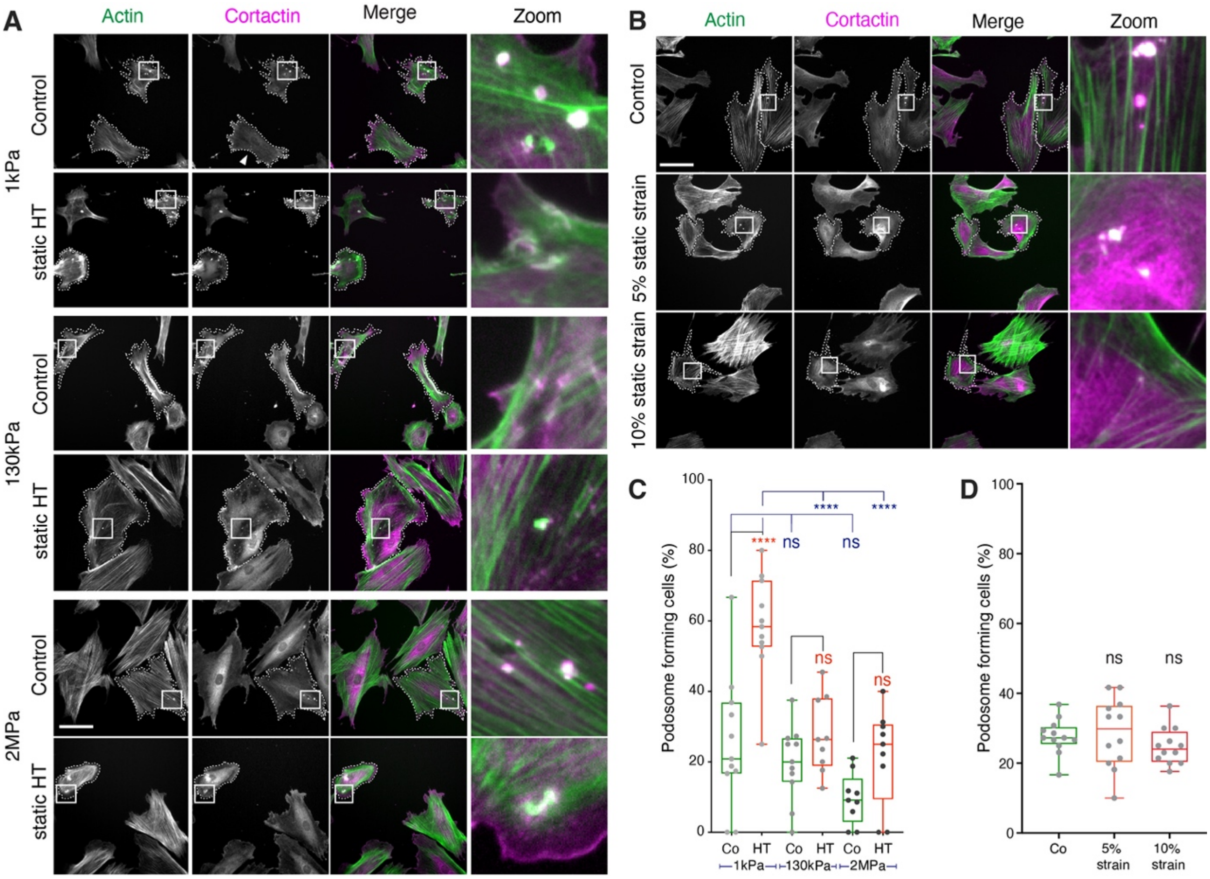


**Supplementary Figure S3: VSMC form podosomes after 30minute static pressure, but not stretch.** A-B) VSMCs were subjected to 30min hydrostatic pressure at 24kPa or alternatively 5% or 10% static biaxial stretch and then stained for Cortactin and Actin. C,D) Quantification of the fraction of podosome forming cells per image (with ~10-20 cells per image), pooled from three independent experiments.

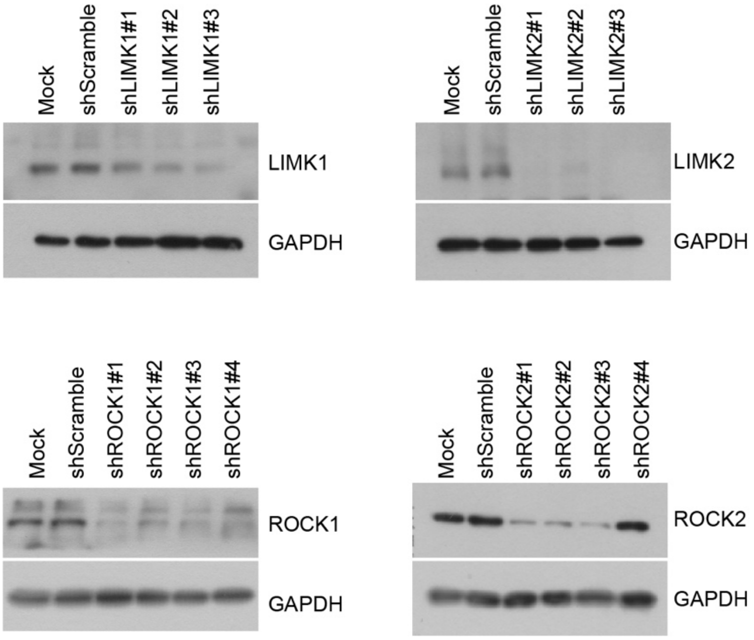


**Supplementary Figure S4: Validation of shRNA knockdowns.** Representative images from three independent repeats.

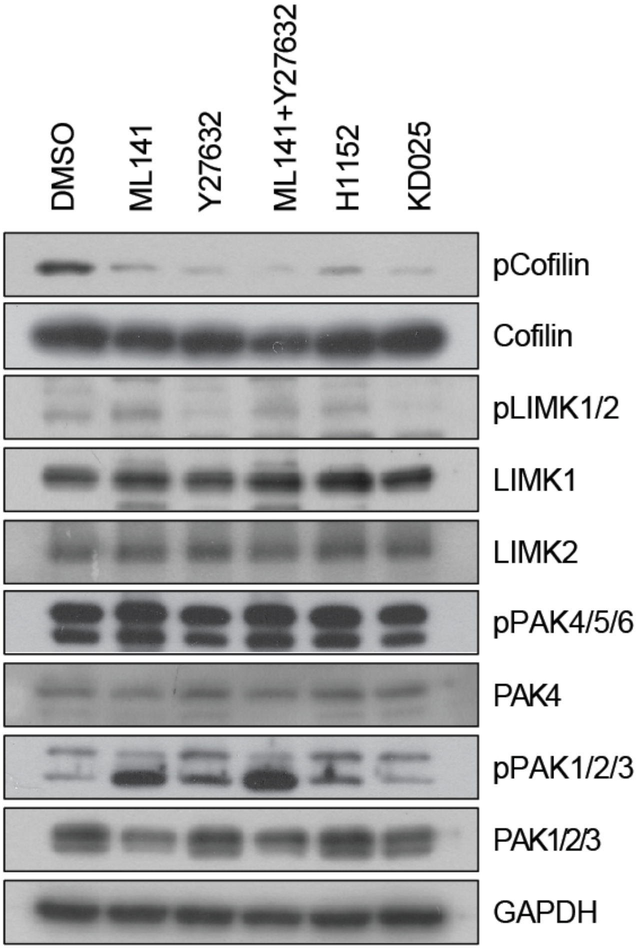


**Supplementary Figure S5: Western blot analysis of effects of Cdc42 (ML141, 10µM), ROCK1/2 (Y27632, 20µM; H1152, 10µM) or ROCK2 (KD025, 10µM) inhibition.** The inhibition of ROCK2 specifically supresses the phosphorylation of both LIMK and cofilin. The inhibition of Cdc42 results in the dephosphorylation of cofilin and the phosphorylation of group 1 PAK (PAK1/2/3) while LIMK phosphorylation remains unchanged. The dual inhibition of ROCK and Cdc42 promotes further cofilin dephosphorylation. All inhibitors are treated for two hours.

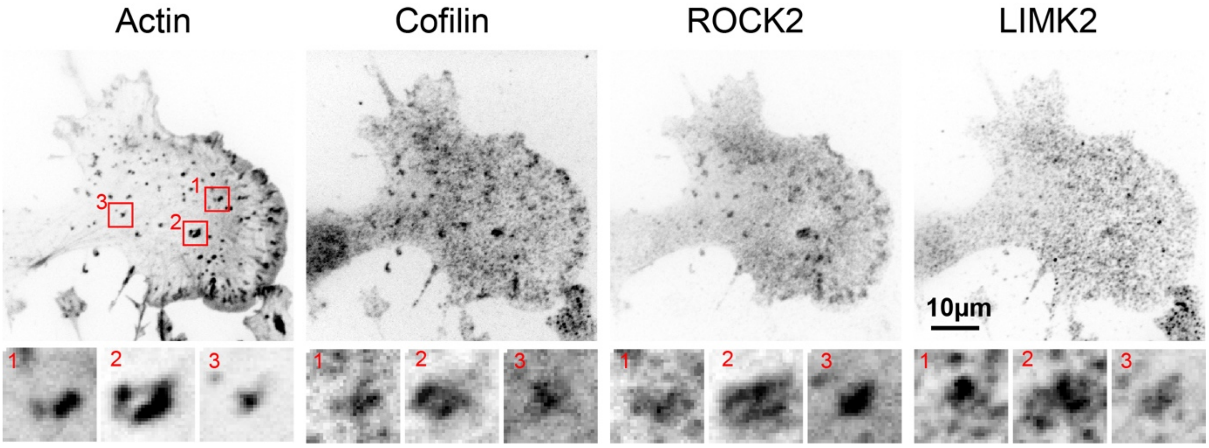


**Supplementary Figure S6: ROCK2 and LIMK2 and cofilin localize to podosomes.** Bottom panel shows zoom of three regions marked in the actin channel.

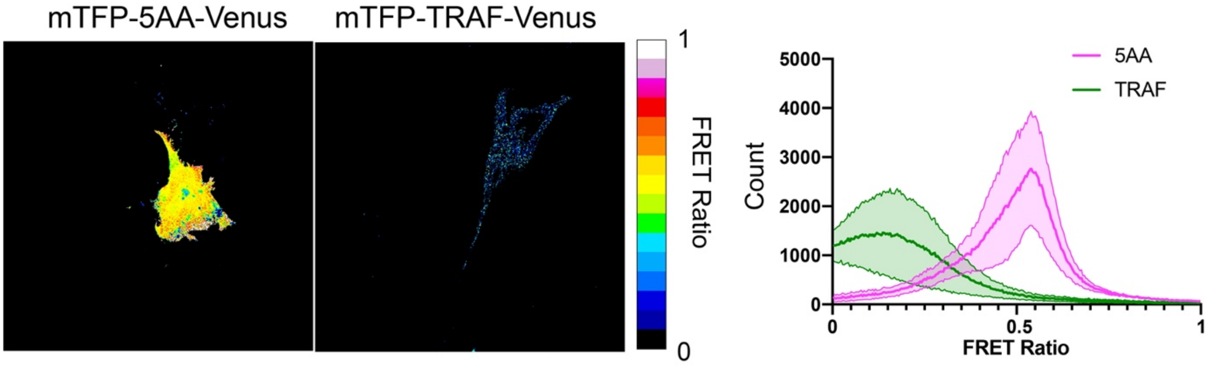


**Supplementary Figure S7: FRET controls.** mTFP-5AA-Venus and mTFP-TRAF-Venus were used to determine the dynamic range of the RhoA-Biosensor on the microscope setup.

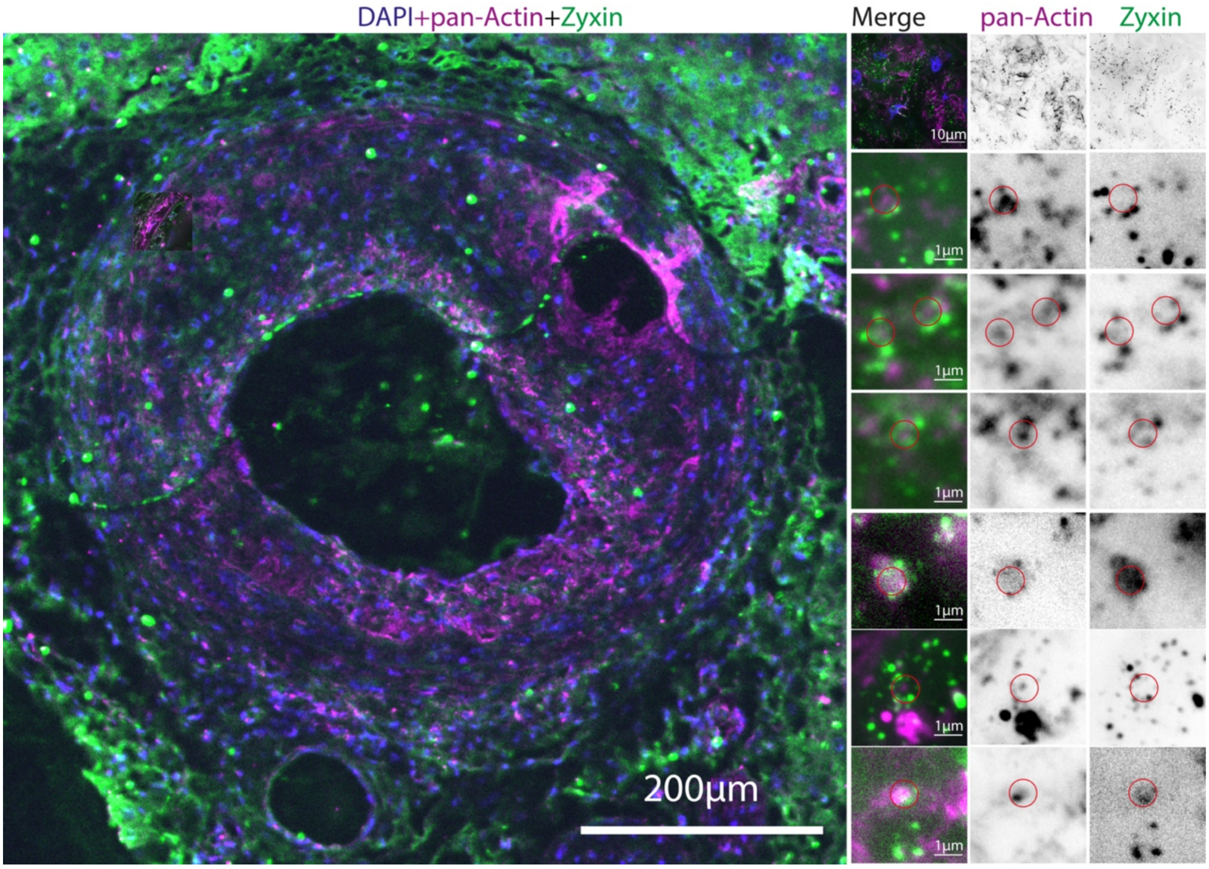


**Supplementary Figure S8: Accumulation of actin and zyxin speckles in neointima.** Left panel shows an overview image of the artery after carotid ligation with clear evidence of neo-intima formation. Top panel on right shows a maximum intensity projection of a z-stack of the neo-intima, taken with a 100x Oil objective on a Nikon SoRa super-resolution spinning disc microscope with 2.8x SoRa magnification. The below panels show further zoom ins with examples of actin dots, surrounded by zyxin areas. Red circles indicate the typical 1µm podosome diameter.

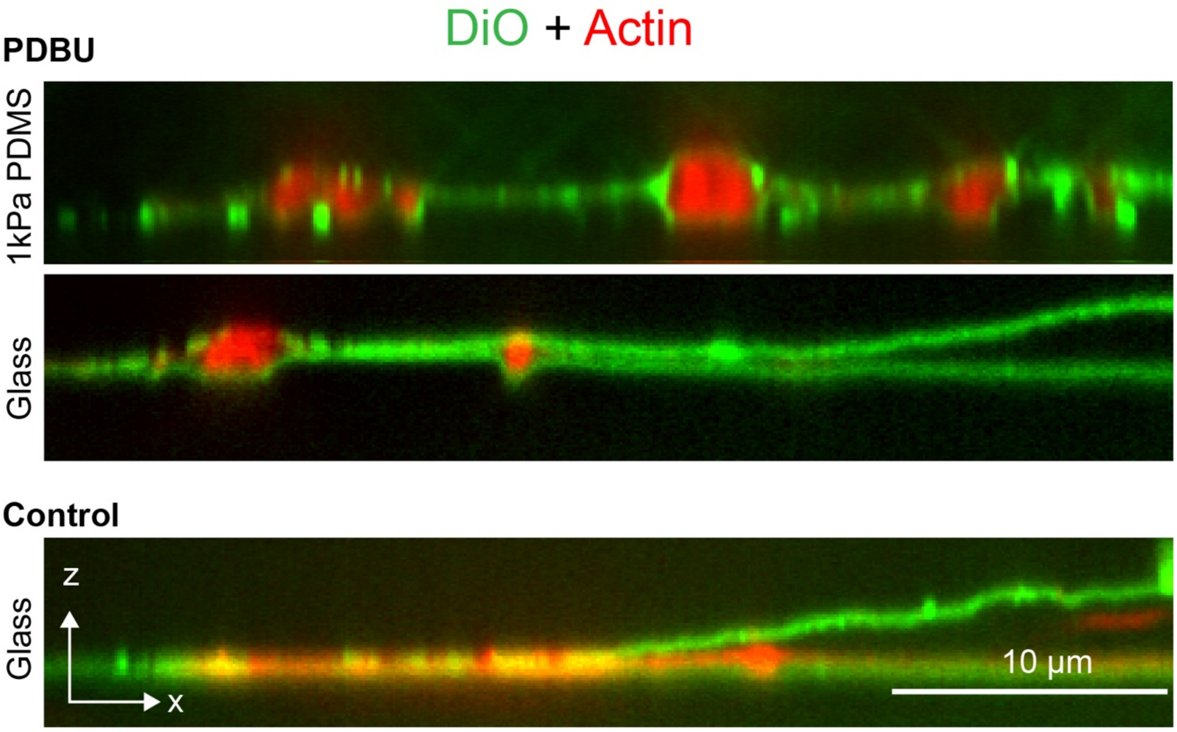


**Supplementary Figure S9: XZ-projection of confocal image stack from VSMCs on Glass or 1kPa PDMS, stained with phalloidin and DiO.** Podosome formation after PDBu treatment results in plasma membrane curving on PDMS and glass, which is absent in control cells with stress fibres.

#### Supplementary Movies:

**Supplementary Movie 1:**
PDBu stimulated vascular smooth muscle cell on 1kPa PDMS, expressing Tractin-Tomato. The images were taken at a rate of 30s per frame.

**Supplementary Movie 2:**
PDBu stimulated vascular smooth muscle cell on 130kPa PDMS, expressing Tractin-Tomato. The images were taken at a rate of 30s per frame.

**Supplementary Movie 3:**
PDBu stimulated vascular smooth muscle cell, expressing a RhoA biosensor and iRFP-Lifeact. Donor and FRET channels were taken simultaneously, using a two camera setup at a rate of 5s per frame. iRFP images were taken every 20 frames (i.e. every 100s).

#### Supplementary Tables

**Supplementary Table 1:**
List of 1027 identified proteins from quantitative proteomic analysis including description, and associated GO terms and pathways.

**Supplementary Table 2:**
Pairwise comparison of quantified proteins from A7r5 VSMCs seeded on 1kPa PDMS and subjected to atmospheric pressure or cyclic HT pressure over 24h.

**Supplementary Table 3:**
Pairwise comparison of quantified proteins from A7r5 VSMCs seeded on 130kPa PDMS and subjected to atmospheric pressure or cyclic HT pressure over 24h.

**Supplementary Table 4:**
Pairwise comparison of quantified proteins from A7r5 VSMCs seeded on 1 or 130kPa PDMS and cultured inside the pressure stimulator at atmospheric pressure over 24h.

**Supplementary Table 5:**
Pairwise comparison of quantified proteins from A7r5 VSMCs seeded on 1 or 130kPa PDMS and subjected to cyclic HT pressure stimulation over 24h.

**Supplementary Note 1:**
The following proteins were included in Figure 2F,G as podosome associated proteins: Calm2(*1*), Macf1(*2*), Arap3(*3*), Pikfyve(*4*), Cgn (included here as interaction partner of ZO-1 in the zona occludens and because of the well described role of the zona occludens complex in podosomes)(*5-8*), Adam6a(*9*), Gbf1(*10*), Rack1(*11*), Myo1e(*12*), Fscn1(*13*), Dstn(ADF)(*14, 15*), Tns1(*16, 17*), TubA1A, Tubb6 (both due to the well-established role of microtubules in podosomal stability regulation)(*18, 19*).

The following proteins were included in Figure 2F,I as atherosclerosis associated proteins:

Vig(*20*), Socs3(*21-23*), Igf2r (*24*), Col6a1(*25*), Col3a1(*25, 26*), Mgp(*27*), Arhgef1(*28*), Stxbp2(*29*)(also part of KEGG set for Atherosclerosis/Lipid), Flnc(*30*), Ttll12(*31*), Rangap1(*32*), RPL30 (part of KEGG set for Atherosclerosis/Lipid), Sema7a(*33*), Fhl1(*34, 35*) Myo1e(*36, 37*).

The following proteins were included in Figure 2F,H as stress response proteins: Hspa8(*38, 39*), Hsp90ab1(*40*), Hspa4(*41*), UBA1(*42, 43*), Ahsa1(*44*), Gstp1(*45*), Psmd3(*46*), Ap3b2(*47*), TubA1A, Tubb6 (both part of Reactome Pathway set Cellular Response to Stress)
